# ORFanes in mitochondrial genomes of marine polychaete *Polydora*

**DOI:** 10.1101/2023.02.04.527105

**Authors:** Maria Selifanova, Oleg Demianchenko, Elizaveta Noskova, Egor Pitikov, Denis Skvortsov, Jana Drozd, Nika Vatolkina, Polina Apel, Ekaterina Kolodyazhnaya, Margarita A. Ezhova, Alexander B. Tzetlin, Tatiana V. Neretina, Dmitry A. Knorre

## Abstract

Most characterised metazoan mitochondrial genomes are compact and encode a small set of proteins that are essential for oxidative phosphorylation. However, in rare cases, invertebrate taxa have additional open reading frames (ORFs) in their mtDNA sequences. Here, we sequenced and analysed the mitochondrial genome of a polychaete worm, *Polydora cf. ciliata*, part of whose life cycle takes place in low-oxygen conditions. In the mitogenome, we found three “ORFane” regions (1063, 427, and 519 bp) that have no resemblance to any standard metazoan mtDNA gene but lack stop codons in one of the reading frames. Similar regions are found in the mitochondrial genomes of three other Polydora species and *Bocardiella hamata*. All five species share the same gene order in their mitogenomes, which differ from that of other known spionidae mitogenomes. By analysing the ORFane sequences, we found that they are under negative selection pressure, contain conservative regions, and harbour predicted transmembrane domains.The codon adaptation indices (CAIs) of the ORFan genes were in the same range of values as the CAI of conventional protein-coding genes in corresponding mitochondrial genomes. Together, this suggests that ORFanes encode functional proteins. We speculate that the ORFanes originated from the conventional mitochondrial protein-coding genes which were duplicated when the Polydora/Bocardiella species complex separated from the rest of the Spionidae.

**Significance statement:** Metazoan mitochondrial genomes usually contain a conservative set of genes and features. However, mitogenomes of some species contain ORFanes – putative protein-coding genes without clear homology with other known sequences. In this study, we analysed three ORFanes in mitochondria of species of the genera Polydora and Bocardiella, which were absent in all other representatives of Spionidae. To the best of our knowledge, ORFanes haven’t been described in Annelida before. Sequence analysis of the ORFanes suggests they contain conservative regions and are likely translated into functional proteins. Our study features an uncommon case where new protein-coding genes emerged in the mitochondrial genomes of metazoa.

## Introduction

Mitochondrial genomes of Metazoa are characterised by the consistency of their genomic features. A typical metazoan mitogenome usually contains 37 genes encoding 13 proteins, 2 ribosomal RNA genes (rRNAs) and 22 transfer RNA genes (tRNAs), as well as a single non-coding ‘control region’ (CR) with regulatory signals for replication and transcription. Such features also include highly conserved genetic architecture and compact gene organisation, which implies the absence of additional non-coding regions, NCRs (Ghiselli et al. 2021). However, some invertebrate species have additional non-coding regions and “unconventional” genes in their mitochondrial genomes (Milani et al. 2013). For instance, the mitochondrial genomes of demosponges harbour an additional ATP-synthase gene, *ATP9* (Lavrov et al. 2005); octocoral mitogenomes contain a DNA-repair mutS gene (Pont-Kingdon et al. 1995). Furthermore, bivalve molluscs have been shown to have additional sex-specific protein-coding genes (Mitchell et al. 2016).

Putative protein-coding genes found in bivalves are called mitochondrial ORFan genes — open reading frames with no detectable homology and unknown function. There is evidence which supports these genes playing a role in mitochondrial DNA inheritance (Mitchell et al. 2016). Mitogenomes of Cnidaria, Porifera and Ctenophora species can also contain ORFans (Flot & Tillier 2007; Breton et al. 2009; Schultz et al. 2020). Some of the ORFans have been shown to have conserved structure across related species (Guerra et al. 2019; Schultz et al. 2020).

Spionids, which make up one of the most diverse families of marine annelids, are characterised by an amazing plasticity of their lifestyles and life cycles. Spionids occur in a wide variety of habitats from the intertidal to the deep sea, sometimes forming dense benthic assemblages (Blake et al. 2020). Their long-living larvae can be the dominant forms of larval plankton in the pelagial. Individuals may extend their palps from burrows or tubes to filter particles from the water; in other cases, the worms are surface deposit feeders and use their palps to sweep the sediment surface. In general, Spionids are characterised by life cycle adaptations such as a wide variety of sperm types and types of fertilisation, asexual reproduction (architomy and paratomy), poecilogony, and in some cases, adelphophagy (Radashevsky & Migotto 2017; Sato-Okoshi et al. 2017). Spionids of genera close to the genus *Polydora* are characterised by an unusual ability to drill hard calcareous substrates. In most cases, they drill into mollusk shells, often being pests of commercially cultivated mollusks (e.g. oysters). At the same time, these animals are often able to build tubes and live in the sediment (Blake et al. 2020).

Recently, three complete mitogenomes of shell-boring Polydora species, *P. websteri, P. brevipalpa*, and *P. hoplura*, were sequenced (Ye et al. 2021; Lee & Lee 2022). Analysis of two related species, *P. websteri* and *P. brevipalpa*, and comparisons to other sedentary animals showed that Polydora mitogenomes have some unusual features: the gene order has been changed, and there are four non-coding regions that are longer than 500 bp. Analysis of four large NCRs in both genomes has shown that none of them exhibited sequence similarity to the nearby coding regions. At the same time, several open reading frames (ORFs) > 50 amino acid residues were detected in NCRs, but BLASTp analysis found no hits with known proteins for any of the putative protein products (Ye et al. 2021). The mitogenome of *P. hoplura* has been shown to have similar architecture, including the presence of NCRs (Lee & Lee 2022).

In this study, we sequenced and assembled the mitogenome of an additional specimen of *Polydora* collected in Biofiltry Bay near White Sea Biological Station. *Polydora ciliata*, originally found on the east coast of Scotland, has a very wide areal (including the North Atlantic and the North Pacific), most likely representing a whole group of very close species (Radashevsky & Pankova 2006, 2013). In the White Sea, it was first recorded about 65 years ago (Sveshnikov 1958). In different parts of this small sea, *P. ciliata* bores into the shells of littoral gastropods, such as *Littorina littorea* on the Solovetsky Islands, or lives in a very rich organic sediment on the border of sulfur-donor (low-oxygen conditions) zones in small shallow inlets (Kandalaksha Bay). We analysed this and previously described *Polydora* and other Spionida species’ mitogenomes and found that three out of four NCR regions lack stop codons along the entire length of the region in one of the reading frames. Moreover, these regions accumulated more synonymous than non-synonymous mutations. This implies that these regions might encode functional proteins.

## Results

We collected a specimen of *Polydora cf ciliata* in the Biofiltry Bay, a small shallow water inlet in the Kandalaksha Bay of the White Sea (see Materials and Methods for details). Isolation and sequencing of total genomic DNA from this specimen enabled assembly of the mitochondrial genome. In this genome, we found all standard metazoan protein-coding genes (PCGs), rRNAs, tRNAs, duplicated tRNAs, as well as four additional regions that were also previously found in *P websteri, P brevipalpa*, and *P. hoplura* species (Ye et al. 2021;Lee& Lee 2022). Similar to other polychaetes, all genes were encoded on a single strand (Figure 1).

**Figure 1.**
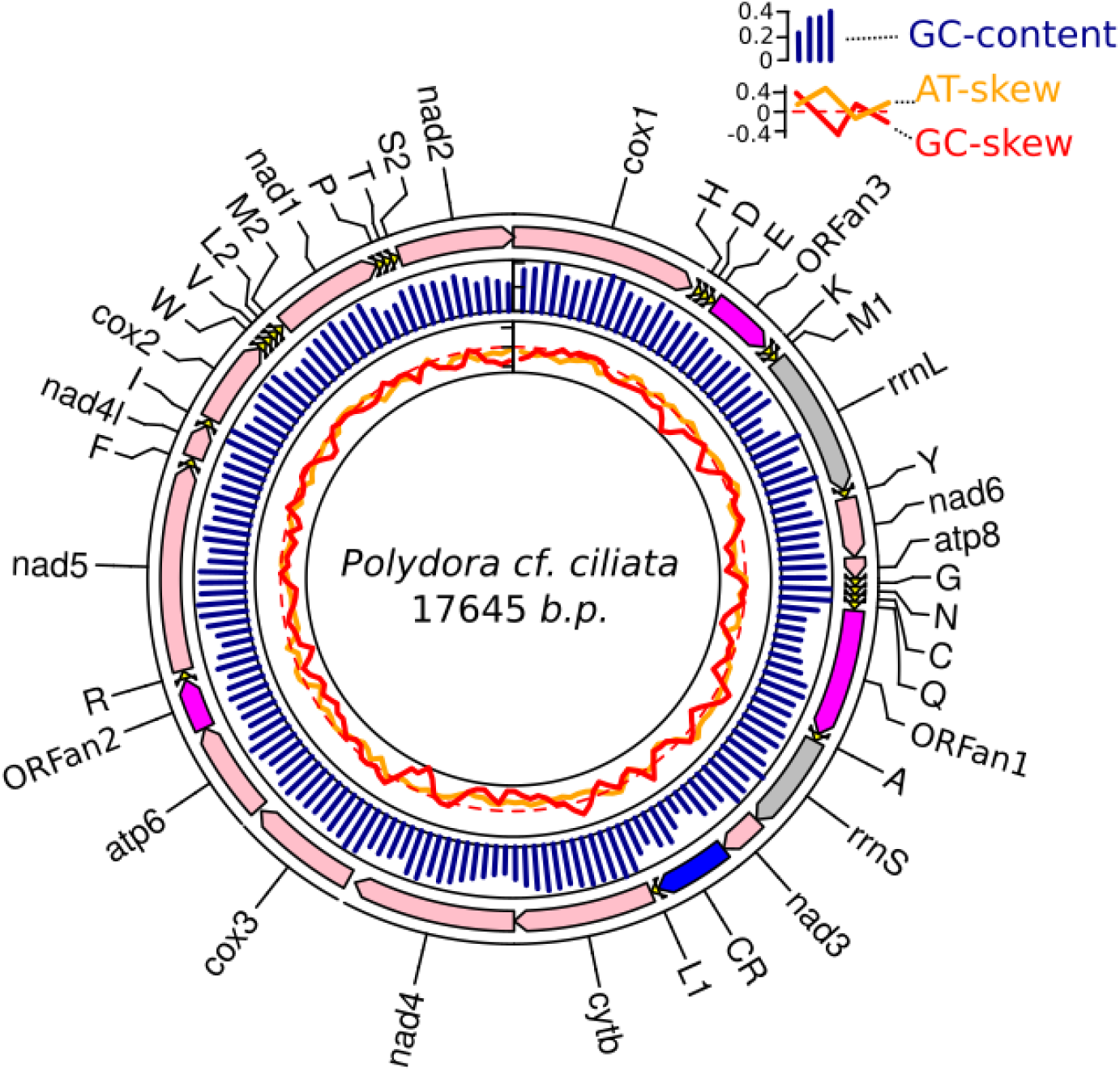
Mitochondrial DNA map of *Polydora cf. ciliata*. Outer tracks represent genes, ORFanes and Control region (CR); GC-content, GC-skew and AT-skew were calculated in 200 bp window (step 100 bp)

Phylogenetic analysis of Spionidae and Polydora mitochondrial genomes available on GeneBank with *Pseudopotamilla reinformis* as an outgroup was made on 12 standard OXPHOS genes. We translated all genes with mitochondrial invertebrate genetic code and aligned the resulting amino acid sequences. We then concatenated alignments and built a phylogenetic tree (Figure 2). The analysis showed that the sequenced mitochondrial genome –*Polydora cf. ciliata* – is close to *P. websteri*. Phylogenetic trees built from codon alignments were consistent with those built from amino acid alignments. To clarify *Polydora cf. ciliata*phylogeny we then built phylogenetic trees from alignments of 16S 18S, and 28S Polydora and Spionidae genes. According to the analysis of 16S rRNA sequences *Polydora cf. ciliata* is closely related to *P. onagowaensis*, although the construction of phylogenies from individual 16S, 18S, and 28S genes did not allow us to obtain reliable topologies Figures S1-S3.

**Figure 2.**
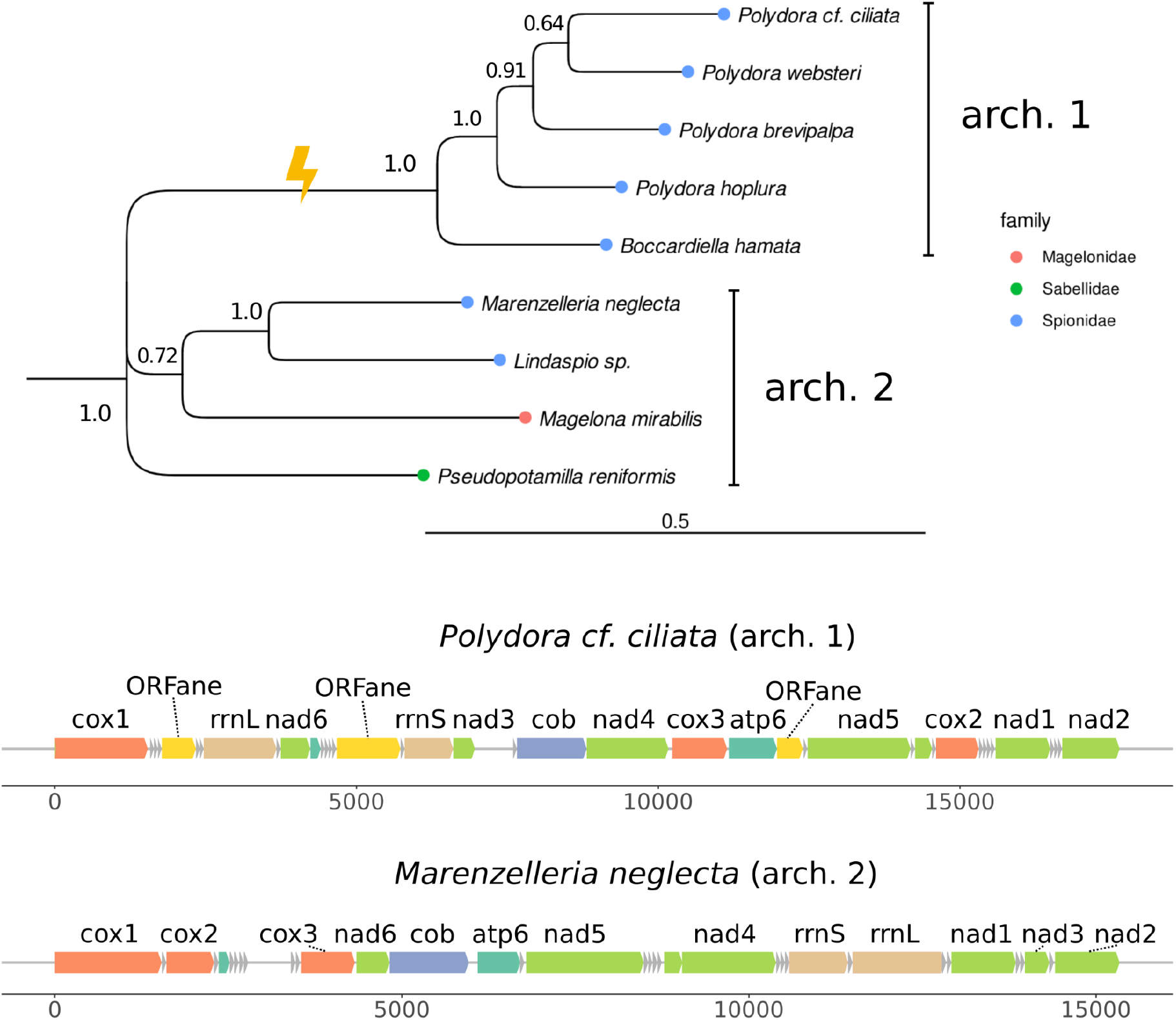
Species tree with genome architecture ORFanes.

Available *Polydora* mitogenomes form a cluster that also includes the *Bocardiella hamata* mitochondrial genome (Figure 2). mtDNAs sequences comprising this cluster harbour four additional regions and have a unique gene order which distinguishes them from the other Spionidae species. To test if the four unconventional regions encode proteins we translated them in all six reading frames with invertebrate mitochondrial genetic code. We found that three of the regions contained one frame without stop codons; this was true in the cases of all four Polydora and *B. hamata* mitogenomes. Figure S4 shows distributions of stop codons in the mitogenomes in all six frames. In three regions all but one frame contained 4-39 stop codons per frame. The fourth region contained stop codons in all frames. Furthermore, this fourth region had lower GC-content than any of protein-coding genes. Finally, it contained sequences capable of folding into stable secondary structure elements, according to the RNAsurface algorithm (see methods and Figures S5). The replication origin predicted by MitoZ (Meng et al. 2019) also falls in this region. Therefore, we annotated it as a control region of the mitochondrial DNA, whereas the other three regions we annotated as ORFanes.

Thereafter, we tried to find sequences with homology to these three unconventional mtDNA regions. First, we tested homology among protein sequences available in databases. A BlastP search of ORFan proteins from *P. cf. ciliata* mitogenome against both NRPD and UniprotKB found no significant hits with e-value cutoff = 1, Tblastx and blastn also found no hits in NCBI databases, while a tblastn search against UniProtKB and NRPD found hits in *Polydora hoplura* mitochondria (e-value: 6e-28) and *Boccardiella hamata* (e-value: 3e-17) mitochondria that are already covered in this study. A PSI-blast search against UniProt found some Bacteria tRNA U34 carboxymethyltransferase hits with an e-value no less than 0.5, which we considered insignificant, and the PSI blast search against NRPD resulted in zero found hits. Thus, we found no detectable homology for any of *P. cf. cliata* ORFan proteins.

In order to detect more remote homologies we also performed an HMM vs sequence search using hmmsearch against available databases (see methods). As a result of the search, no significant hits (with the cutoff value of 0.01) were found in any of the databases for both the first and the second ORFan proteins. For the third ORFan protein, hmmsearch found one significant hit (e-value=0.0028) in the ‘reference proteomes’ database and two significant hits in UniProtKB (e-values=0.0009 and 0.0083). Tertiary structure predictions and structure similarity searches for three ORFanes in five species were performed using @tome and i-tasser. More detailed reports on @tome and i-tasser results are shown in Table S3, Text S2. The inconsistencies in the taxonomy and functions of the found proteins as well as relatively weak scores of the hits made for ORFans of different species do not allow us to draw conclusions about the significance of the detected putative homologies.

Meanwhile, the alignment of the ORFanes from *Polydora cf. ciliata* mitogenome to the corresponding ORFanes in other mitogenomes revealed that all of them have conserved regions (Figure 3). To obtain a conservation score track, we build alignments of three ORFanes as well as two conventional protein-coding genes, *NAD6* and *CO1*, in all five species of consideration. We used MEGA and jalview by Muscle with the default algorithm (see methods for details). In ORFanes, both regions with high and low conservation scores were found. For the ORFane-1, the most conservative regions were in the positions: 213-240, 261-282; ORFane-2, 7-15 and 32-72 regions were the most conservative. ORFane-3 had the largest number of regions with high conservation scores: 1-14, 45-56, 71-97, 119-180.

**Figure 3.**
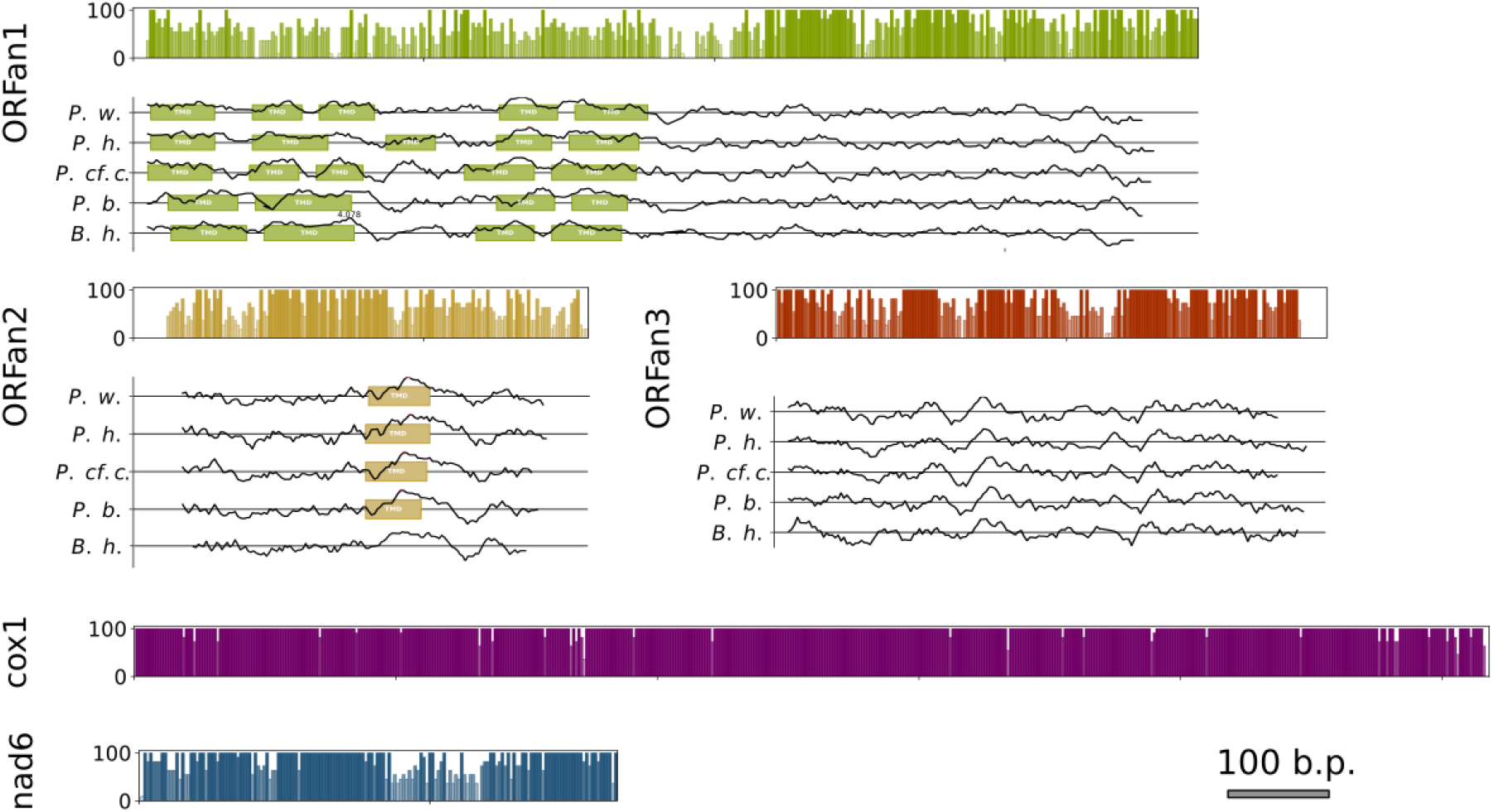
ORFanes contain conservative regions and transmembrane domains. Conservation, domain architecture, and hydrophobicity profiles for ORFans 1-3, *NAD6* and *CO1* genes. The transmembrane regions were predicted using Phobius and TMHMM, hydropathy profiles were calculated with the method of Kyte and Doolittle, ProtScale tool at ExPASy. *P.w. – Polydora websteri; P.h. – Polydora hoplura; P. cf. c. – Polydora cf ciliata; P.b. – Polydora brevipalpa; B.h. – Bocardiella hamata*

To determine the possible function and structure of ORFanes, we predicted the domain architectures of ORFanes and made hydrophobicity tracks in five studied species. We predicted transmembrane domains using Phobius integrated into InterProScan (see Materials and Methods). Phobius predicted the presence of several transmembrane domains in ORFanes 1 in the N-terminus part of the putative protein in all species, whereas ORFanes 2 harboured a single transmembrane domain located closer to the C-terminus of the protein in the same position in all analysed species. Transmembrane domains have been predicted for ORFans 1 and 2 and not for ORFan 3 (Figure 3). It is important to note that algorithms integrated into InterProScan didn’t find any functional domains in ORFan proteins other than transmembrane domains. In order to address the possible function of the proteins, we also searched for functional motifs in the sequences using MotifScan. Only two hits were marked by MotifScan as “strong” while others were either “weak” or “questionable”. In the ORFan1 from *Polydora brevipalpa*, MotifScan found a Phenylalanine-rich region profile (positions 8-170), and in the ORFan1 from *Polydora hoplura*, it found a Leucine-rich region (2-154) and a Phenylalanine-rich region (6-47). Both TPRpred and SignalP didn’t find any Peptide Repeats or Signal Peptides respectively in Polydora ORFanes.

The absence of stop codons in one of the reading frames of the ORFanes suggests that they may code for proteins. In this case, one would expect negative selection pressure on their protein sequence, which should be expressed in an increased dn/ds ratio. To test this, we calculated nucleotide and amino acid p-distances as well as the dn/ds ratio for protein-coding genes in *Polydora* and *Bocardiella* genomes (Figure 4). ORFane genes appeared to be the most diverse of the analysed genomes (Figure 4A, B). Thus, we consider them to be the fastest-evolving protein-coding genes in the mitogenomes. Accordingly, as shown in Figure 4C, dn/ds ratios in ORFanes are also slightly higher than in other protein-coding genes. However, the codon-based Z-test of purifying selection showed that along with the other protein-coding genes, ORFans are subject to negative selection in all genomes investigated, which is statistically significant (p-value less than 0.001). This finding indicates that ORFans are likely translated to the functional proteins and are the subject of negative selection. Furthermore, we calculated Codon Adaptation Indexes (see methods) for PCG and ORFanes, using information from all other genes as a background. CAIs reflect codon usage bias and are expected to be high in weakly expressed genes. Figure 4D shows that the CAI of ORFan genes is similar to the CAI of other protein-coding genes; in some worm mitogenomes it was even higher than the CAIs of some conventional PCGs. e.g. *NAD3* and *NAD4l*.

**Figure 4.**
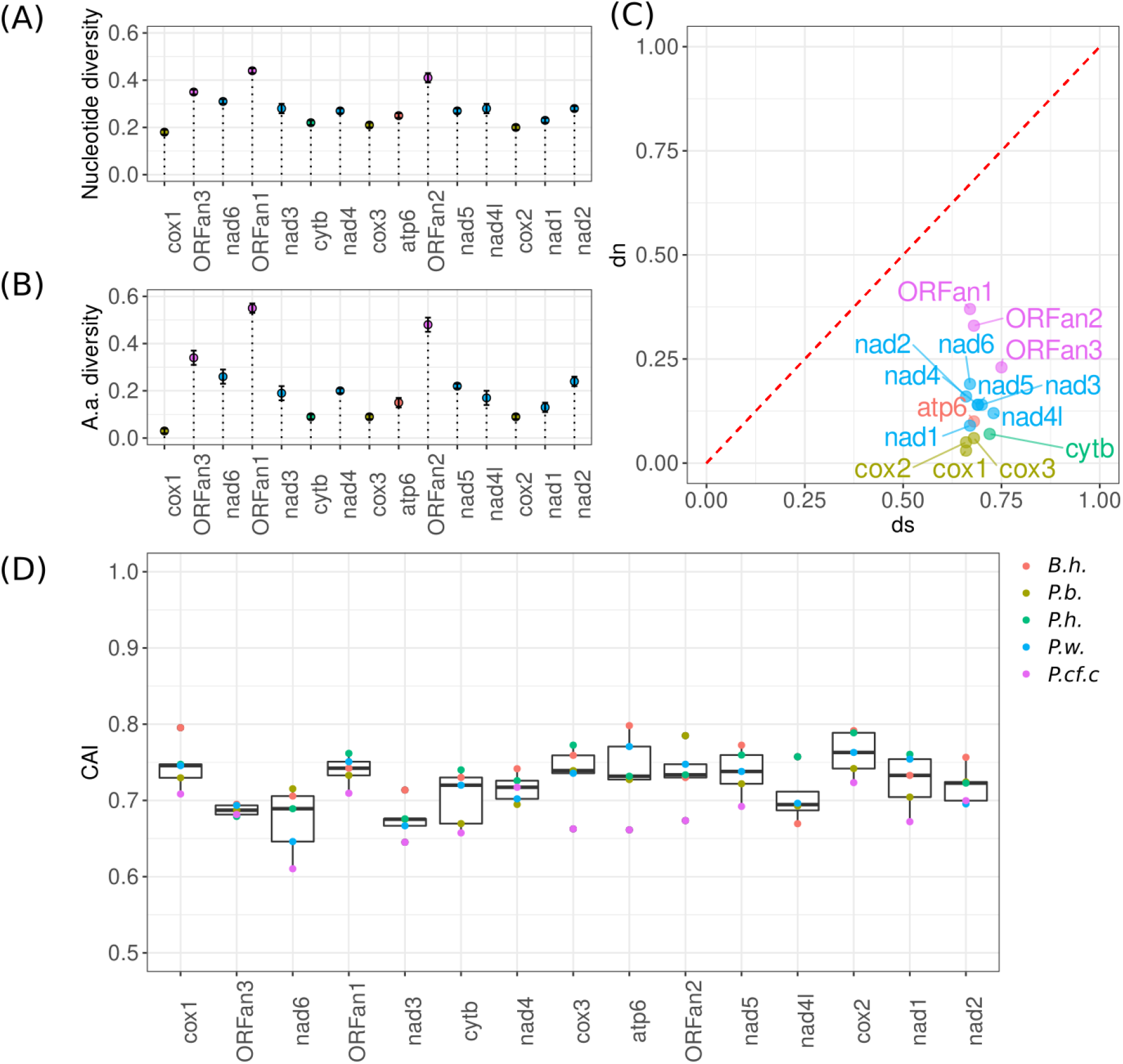
ORFanes evolve faster than conventional PCGs but are under the pressure of negative selection. Nucleotide (A) and amino acid (B) diversity of the ORFanes and other mitochondrial PCGs. Values for variance of nucleotide and amino acid p-distances are obtained by the bootstrap method with 500 replicates. (C) Proportion of synonymous (ds) and non-synonymous mutations. The codon-based Z-test for purifying selection shows that we should reject the null hypothesis of neutral evolution (dn=dn) in favour of the alternative hypothesis of purifying selection for all standard OXPHOS genes as well as for all ORFan genes (p-value << 0.05). (D) Codon adaptation index of mitochondrial PCGs and ORFanes.

## Discussion

Mitochondria evolved from a proteobacteria endosymbiont (Martijn et al. 2018). After symbiogenesis, its genome has undergone a significant reduction. Many genes have been lost or transferred to the nuclear genome while acquisitions of new elements in mtDNA via horizontal transfer were extremely rare (Janouškovec et al. 2017). One of the exceptions is the mutS gene in the mitochondrial genomes of octocorals (Pont-Kingdon et al. 1998). Gene duplications within mitochondrial genomes are also uncommon, although reported in several taxa. For instance, mitogenomes of the nematode *Caenorhabditis briggsae* contain pseudogenized *NAD5* (Raboin et al. 2010). The mitochondrial genome of *Chaetopterus variopedatus*, a parchment worm, contains a duplicated *COX1* gene and a portion of a duplicated *NAD3* gene. The latter is adjacent to an unannotated region (Weigert et al. 2016). In this study, we expand these rare examples by the ORFanes located in the mitochondrial genomes of Polychaete worms in the genus *Polydora*.

Mitochondrial ORFanes can be of three possible origins: (1) they were present in the mitochondrial genome of the metazoan’s common ancestor and preserved in some clades; (2) they arose through duplication of conventional mtDNA genes; or (3) they emerged via horizontal transfer from other genomes. The first explanation is highly unlikely given that all other sequenced annelid mitogenomes are devoid of ORFanes, but there is not enough data to discriminate between the second and third possibilities. On the one hand, ORFanes have no homology with any other mitochondrial protein-coding gene. On the other hand, if ORFanes have recently been transferred into the mitochondrial genome from a virus or any nuclear genome, we would expect significant differences in AT-content, AT-skew and GC-skew. Figure 1 and Figure S5 show that these genomic parameters of ORFanes fall in range of the same parameters of conventional PCGs.

We suggest that ORFanes emerged as a result of intrachromosomal rearrangement of the *Polydora* ancestor and then underwent a stage of rapid evolution that destroyed the phylogenetic signal that could allow us to identify their origin. Indeed, tandem duplications in mitochondrial genomes are usually followed by random or non-random gene loss. These events result in rearrangements in mitochondrial DNAs (Lavrov et al. 2002; San Mauro et al.2006). At the same time, one of the duplicated genes can undergo rapid evolution and neofunctionalisation (Conant & Wolfe 2008). We suggest that the mitochondrial genome of a common ancestor of *Polydora* and *Bocardiella* worms was duplicated and/or reshuffled. After that, duplicated genes were either lost or repurposed in their role and function. The fact that genomic rearrangement and ORFane emergence are on the same branch of the phylogenetic tree (Figure 2) votes for this otherwise speculative assumption.

Polydora ORFanes are unusual from two points of view. First, mitochondrial genomes of other sequenced annelida, a large phylum, are devoid of uncommon PCGs. Second, genomic data suggests that these genes are transcribed and translated into the functional proteins. All ORFanes in all five analysed species are devoid of stop codons (Figure S4). Moreover, some regions of ORFanes show a high degree of conservation, and ORFane 2 contains a predicted transmembrane domain in all five analysed species in a similar genomic position (Figure 3). Finally, ORFane sequences showed evidence of negative selection: kn/ks are 0.552 (ORFane1), 0.485 (ORFane2), 0.307 (ORFane3) (Figure 3C). It should be mentioned, that these analyses cannot exclude the possibility that ORFanes recently became non-functional (useless to worm mitochondria) in some of the studied species and did not accumulate enough mutations to detect them. However, the scenario suggesting that ORFanes became pseudogenes in all five species is highly unlikely.

In male-transmitted mtDNA mitochondrial genomes of bivalves, some standard PCGs are significantly elongated, whereas additional sequences show no homology to any other known sequence (Ghiselli et al. 2021). As a result, the size of the corresponding protein is significantly increased (Tassé et al. 2022). It is noteworthy that all three ORFane genes are in frame with previous PCGs (Figure S4), although ORFanes 1 and 3 are separated from the upstream proteins by multiple stop codons and tRNAs (Figure 1). Meanwhile, there is only one stop codon between *ATP6* and ORFane 2. At the same time there is a possibility of translational readthrough — the ability of the ribosome to bypass stop codons (Mangkalaphiban et al. 2021). In yeast, nuclear genome translational readthrough can reach up to 10% of the total proteins (Fearon et al. 1994). Thus, we cannot exclude that ORFane 2 translated as an extension of the ATP6-encoded ATP-synthase subunit *a* under some specific conditions. Conversely, it might be just a consequence of invertebrate mtDNA being transcribed into and translated as polycistronic mRNAs.

To summarise, in this study we analysed three putative protein coding genes in mitochondrial genomes of several polychaete species belonging to the genus *Polydora* and *Bocardiella*. These genes are likely to encode proteins with conservative amino-acid sequences, although we detected no homologues of the ORFanes outside of this group. Moreover, their possible functional role cannot be deduced from their sequences, although it might be noted that all species with ORFanes in mitogenomes rely on drilling of hard calcareous substrates. That might require mitochondria in some tissues to provide extra energy necessary to penetrate calcareous matter. Nonetheless, ORFanes in the *Polydora* and *Bocardiella* species provide a very rare example where new genes emerged in the mitochondrial genome and likely acquired a new function.

## Materials and methods

### Specimen collection and identification

Live specimens of *Polydora cf. ciliata* (Johnston, 1838) (Jacobi, 1883) were collected with the help of SCUBA divers on the soft bottom in the Biofiltry Bay a small shallow water inlet in the Kandalaksha Bay of the White Sea at a depth of 7 - 8 (66° 32’ 21”N, 33° 09’59 “E), in close vicinities of the White Sea Biological Station of the MSU.

Spionid polychaetes *Polydora ciliata* were first described by Johnston in 1838 (Johnston 1838) as inhabitants of muddy tubes in shallow waters (littoral) on the coast of Scotland (Berwick Bay). Since then worms very similar in morphology to those described by Johnston have been found in both the North Atlantic and the North Pacific. Representatives of the species are described as mass inhabitants of soft bottom substrates, living in silty pipes, or as worms boring holes in mollusk shells (see (Mustaquim 1986; Radashevsky & Pankova 2006) for review). At different times, the inhabitants of silty tubes and bored shells were described as separate but hardly recognizable morphological species (Annenkova 1934); apparently, there is a group of very close morphological species here. However, despite many years of efforts (Kendall 1980; Mustaquim 1986; Manchenko & Radashevsky 1993; Radashevsky & Pankova 2006) this issue still remains unresolved, and it seems to us correct to define our worms as *Polydora cf.ciliata* (Johnston, 1838).

### DNA isolation and NGS sequencing

DNA was extracted using a Diatom DNA kit (Isogen) according to the manufacturer’s recommendations. DNA libraries were constructed using the NEBNext Ultra II DNA Library Prep Kit by New England Biolabs (NEB) and the NEBNext Multiplex Oligos for Illumina (Index Primers Set 1) by NEB following the manufacturer’s protocol. The samples were amplified using 10 cycles of PCR. The constructed libraries were sequenced on an Illumina MiniSeq with paired-end read length of 150.

### Assembly and annotation

The quality of the library was assessed using the fastqc program (Andrews 2010). Primary processing of readings (trimming) was carried out using the trimmomatic program (Bolger et al. 2014). The final assembly was made using SPAdes (Antipov et al. 2019) and the longest contig containing several mitochondrial genes was defined as mitochondrial. The assembly was then refined using Sanger sequencing of unannotated regions and the cox3 gene as well as Illumina sequencing of mitocontig parts obtained using long-range PCR (the primers are listed in Tables S1, S2)

Annotation of the genome was retrieved using MitoZ (Meng et al. 2019). We manually specified the coordinates of PGCs by aligning the genes with the corresponding genes of other annelid species. The control region was identified using the RNASurface algorithm (Soldatov et al. 2014) which reflects the significance of putative secondary structure in the region, as well as by examining GC content distribution along the genome and locating replication origin using MitoZ, Figure S5. The atp8 gene was identified in all 5 considered mitogenomes using multiple alignments of atp8 genes from related species with a potential atp8 ORF, Figure S6.

In order to find ORFanes in all genomes considered we first divided genomes into codons in all reading frames and identified invertebrate mitochondrial stop codons. Then pairwise distances between neighbouring stop codons were calculated and the resulting genomic regions were filtered by length (360 bp). Long regions were divided by t-RNA genes boundaries (if intersected). The resulting possible protein-coding genes were filtered by length again (210 bp). All steps were performed using R programming language (dplyr *u*biostrings libraries). Codon alignments were used to establish exact ORFane (protein-coding genes in unannotated regions) boundaries.

A program code for constructing a schematic image of the mentioned maps and annotations was written in R using circlize library (Gu et al. 2014) and tidyverse libraries (Wickham et al. 2019). Sequence statistics for the plot were computed with Biostrings (Pagès et al. 2017)and dplyr (Wickham et al. 2022). The mapping was performed using bowtie2 (Langmead & Salzberg 2012), and base-wise coverage was retrieved with samtools (Danecek et al. 2021) and computed with dplyr.

### Sequence analysis and phylogeny

Nucleotide and amino acid alignments of standard protein-coding genes and ORFanes were performed using the Muscle algorithm with default parameters (Edgar 2004) integrated in MEGA11 software (Tamura et al. 2021). Tracks of position conservation were obtained with jalview 2.11.2 (Waterhouse et al. 2009). In order to perform a phylogenetic analysis of the studied mitochondrial genomes we then performed the 12 codon and amino acid alignments of standard protein-coding genes with the corresponding genes from Spionidae mitogenomes available on GeneBank and used *Pseudopotamilla reinformis* as an outgroup. A total of 9 mt genomes was therefore considered for the phylogenies. The Atp8 gene was excluded from the analysis because of its high variability and short length. Codon alignments used to construct phylogenetic trees were computed using the altorythm integrated in MACSE tool (Ranwez et al. 2011). Both protein and codon alignments were then merged and used to build phylogenetic trees. In order to clarify the phylogenetic position of Polydora collected in Biofiltry Bay we also constructed phylogenetic trees of the 16S, 18S and 28S genes of different *Polydora* species available on GeneBank (Figures S1-S3). All trees were constructed using the Maximum Likelihood Algorithm integrated in MEGA11 software with default parameters, 100 bootstrap replicas and General Time Reversible model.

Nucleotide and amino acid p-distances, as well as dn/ds ratios and the codon-based test of purifying selection using the Nei-Gojobori method (Nei & Gojobori 1986), were calculated using MEGA11 (Tamura et al. 2021). Sequence comparison plots were built with the ggplot (Wickham 2016) package in R. The codon adaptation index for protein-coding genes was calculated using the Python CAI module (Python Implementation of Codon Adaptation Index) (Lee 2018) with all protein-coding genes of the considered mitogenome set as background.

### Functional analyses of ORFans

For ORFanes, cox1 and nad6 gene domain architectures were obtained using InterProScan 5.59-91.0 (Jones et al. 2014). Hydropathy profiles of amino acid sequences were calculated with the ProtScale tool at ExPASy (Gasteiger et al. 2005), by the method of Kyte and Doolittle (Kyte & Doolittle 1982).

In order to predict possible origins and functions of ORFane proteins we performed a sequence search with several search algorithms with default parameters: blastn, tblastn, tblastx against NCBI databases, blastp and PSI-blast [National Center for Biotechnology Information (NCBI)[Internet]. Bethesda (MD): National Library of Medicine (US), National Center for Biotechnology Information; [1988] – [cited 2017 Apr 06]. Available from: https://www.ncbi.nlm.nih.gov/] against either entire non-redundant protein database (NRPD) and UniprotKB (Release 2022_04).

As suggested in (Mitchell et al. 2016), in order to detect more remote possible homologies we used methods based on profile hidden Markov models (profile HMMs). We used hmmsearch 3.3.2 (Potter et al. 2018) to perform a profile HMM - protein sequence comparison with default parameters against Reference Proteomes, UniProtKB, SwissProt, PDB and AlphaFold databases, where ORFanes’ alignments were used as an input. We also performed a profile HMM – profile HMM comparison against default databases representing proteins with known structure via the latest version of HHpred with default parameters (Zimmermann et al. 2018).

The tertiary structure prediction and the search for structural similarity were performed using @tome v3 (Pons & Labesse 2009) and I-tasser 5.1 (Zheng et al. 2021). In order to address possible functions of ORFan proteins we searched for known motifs that occur in sequences using MotifScan against default databases (https://myhits.sib.swiss/cgi-bin/motif_scan). Sequences were also searched for Tetratrico Peptide Repeats (TPRs) and Pentatrico Peptide Repeats (PPRs) using TPRpred 11.0 (https://toolkit.tuebingen.mpg.de/tools/tprpred). We checked for possible presence of Signal Peptides in ORFan proteins using SignalP 3.0 (Bendtsen et al. 2004).

## Acknowledgements

We are very grateful to Anna Sokolova for her help in editing the grammar and style of the text.

## Data Availability Statement

*Polydora cf. ciliata* mitogenome is submitted to genbank, accession number: OQ078742

## Authors contribution

AT obtained and identified the specimen; ME and TN isolated DNA, prepared the libraries, performed NGS sequencing and verified ORFanes with Sanger sequencing; MS, OD, EN, EP, DS, PA, JD, EK and NV performed computational analysis and prepared the illustrations. MS, AT, DK drafted manuscript text; MS, TN and DK supervised the project; TN obtained funding; All authors participated in conceptualisation, manuscript writing and approved the final version of the manuscript. The contribution of the authors is described in more detail in Supplementary materials (Text S2).

## Supplementary materials

**Table S1.** Long range PCR primers. These primers were used to enrich mitochondrial DNAs, PCR products were sequenced by Illumina, reads were mapped to the assembly and used to verify it.

**Table S2.** Primers flanking unannotated regions. These primers were used to confirm sequences of ORFanes and CR using Sanger sequencing.

**Table S3.** Atome3 and I-tasser top 10 hits (for Atome3 score threshold of 40.00 also applied, for I-tasser duplicate hits for one protein are excluded) of the mitochondrial Polydora and Boccardiella species ORFans.

**Figure S1.** Phylogenetic tree of 16S sequences of Polydora cf. ciliate and related Annelida species constructed using the Maximum Likelihood Algorithm with General Time Reversible model integrated in MEGA11 software with default parameters and 100 bootstrap replicas.

**Figure S2.** Phylogenetic tree of 18S sequences of Polydora cf. ciliate and related Annelida species constructed using the Maximum Likelihood Algorithm with a General Time Reversible model integrated in MEGA11 software with default parameters and 100 bootstrap replicas.

**Figure S3.** Phylogenetic tree of 26S sequences of Polydora cf. ciliate and related Annelida species constructed using the Maximum Likelihood Algorithm with a General Time Reversible model integrated in MEGA11 software with default parameters and 100 bootstrap replicas.

**Figure S4.** Maps of all analysed mitogenomes with visualised stop codons in all six frames. Outer circle shows annotation for a particular genome, three mid-circles show the amount of stop codons per bin for each frame, where red indicates stop codons in forward strand, black in reverse strand. Inner circle shows potential protein-coding regions after tRNA-punctuation of long regions lacking stop-codons (see methods). Species transcript: *P. cf. c. – Polydora cf. ciliata, P. h. – Polydora hoplura, P. w. – Polydora websteri, P b. – Polydora brevpalpa, B. h. – Boccardiella hamata*.

**Figure S5.** Figure S5. A. Analysis of the nucleotide content of Polydora and Bocardiella mitochondrial genes and putative control region (CR) (GC-content, A-T and G-C skew) B. Analysis of the nucleotide sequence for its ability to form secondary structures in a single-stranded state. We analysed three ORFan genes, tRNA-Gly, CO1 and a possible control region. The ability of a sequence to form a stable secondary structure was measured by computing Z-scores; higher values of square minimum Z-score reflect higher significance of secondary structure in the region. The possible presence of regulatory secondary structures was addressed using RNASurface server.

**Figure S6.** Multiple alignments of ATP8 genes. Given the highly conservative nature of mitogenome architecture in Polydora genus and *Bocardiella hamata* we searched for atp8 right after nad6 in the same position where the gene was located in species with annotated atp8. We performed multiple alignment of atp8 genes from Polydora and Bocardiella species [Muscle with defaults parameters, JalView] (A). The sequences proved to be nearly identical, thus we concluded that these ORFs are indeed atp8 sequences not found by automatic annotation. We conducted multiple alignment of all known sedentarian atp8 sequences [using Muscle algorithm with defaults parameters] (B). The alignment revealed substitution of the conserved part of the protein: the first four residues consensus appears to be MPHL instead of MPQL. Although we see the same three first a.a. residues in Polydora genus, its atp8 exhibits exceptional differences from other sedentarian species.

## References

Andrews S. 2010. A quality control tool for high throughput sequence data. https://www.bioinformatics.babraham.ac.uk/projects/fastqc/.

Annenkova N. 1934. Kurze 0bersicht der Polychaeten der Litoralzone der Bering-insel (Kommandor-lnseln), nebst Beschreibung neuer Arten. Zool. Anz. 106:322–331.

Antipov D, Raiko M, Lapidus A, Pevzner PA. 2019. Plasmid detection and assembly in genomic and metagenomic data sets. Genome Res. 29:961–968.

Bendtsen JD, Nielsen H, von Heijne G, Brunak S. 2004. Improved prediction of signal peptides: SignalP 3.0. J. Mol. Biol. 340:783–795.

Blake JA, Maciolek NJ, Meißner K. 2020. Pleistoannelida, Sedentaria II. De Gruyter.

Bolger AM, Lohse M, Usadel B. 2014. Trimmomatic: a flexible trimmer for Illumina sequence data. Bioinformatics. 30:2114–2120.

Breton S et al. 2009. Comparative mitochondrial genomics of freshwater mussels (Bivalvia:Unionoida) with doubly uniparental inheritance of mtDNA: gender-specific open reading frames and putative origins of replication. Genetics. 183:1575–1589.

Conant GC, Wolfe KH. 2008. Turning a hobby into a job: how duplicated genes find new functions. Nat. Rev. Genet. 9:938–950.

Danecek P et al. 2021. Twelve years of SAMtools and BCFtools. Gigascience. 10. doi:10.1093/gigascience/giab008.

Edgar RC. 2004. MUSCLE: a multiple sequence alignment method with reduced time and space complexity. BMC Bioinformatics. 5:113.

Fearon K, McClendon V, Bonetti B, Bedwell DM. 1994. Premature translation termination mutations are efficiently suppressed in a highly conserved region of yeast Ste6p, a member of the ATP-binding cassette (ABC) transporter family. J. Biol. Chem. 269:17802–17808.

Flot J-F, Tillier S. 2007. The mitochondrial genome of Pocillopora (Cnidaria: Scleractinia)contains two variable regions: the putative D-loop and a novel ORF of unknown function. Gene. 401:80–87.

Gasteiger E et al. 2005. Protein Identification and Analysis Tools on the ExPASy Server. In:The Proteomics Protocols Handbook. Walker, JM, editor. Humana Press: Totowa, NJ pp.571–607.

Ghiselli F et al. 2021. Molluscan mitochondrial genomes break the rules. Philos. Trans. R.Soc. Lond. B Biol. Sci. 376:20200159.

Guerra D et al. 2019. Variability of mitochondrial ORFans hints at possible differences in the system of doubly uniparental inheritance of mitochondria among families of freshwater mussels (Bivalvia: Unionida). BMC Evol. Biol. 19:229.

Gu Z, Gu L, Eils R, Schlesner M, Brors B. 2014. circlize Implements and enhances circular visualization in R. Bioinformatics. 30:2811–2812.

Janouškovec J et al. 2017. A New Lineage of Eukaryotes Illuminates Early Mitochondrial Genome Reduction. Curr. Biol. 27:3717–3724.e5.

Johnston G. 1838. Miscellanea Zoologica. Aricidae. Magazine of Jzoology and Botany,Edinburgh. 2:63–73.

Jones P et al. 2014. InterProScan 5: genome-scale protein function classification. Bioinformatics. 30:1236–1240.

Kendall MA. 1980. Variations in some morphological characteristics of Polydora ciliata(Johnston). J. Nat. Hist. 14:405–411.

Kyte J, Doolittle RF. 1982. A simple method for displaying the hydropathic character of aprotein. J. Mol. Biol. 157:105–132.

Langmead B, Salzberg SL. 2012. Fast gapped-read alignment with Bowtie 2. Nat. Methods. 9:357–359.

Lavrov DV, Boore JL, Brown WM. 2002. Complete mtDNA sequences of two millipedes suggest a new model for mitochondrial gene rearrangements: duplication and nonrandom loss. Mol. Biol. Evol. 19:163–169.

Lavrov DV, Forget L, Kelly M, Lang BF. 2005. Mitochondrial genomes of two demosponges provide insights into an early stage of animal evolution. Mol. Biol. Evol. 22:1231–1239.

Lee BD. 2018. Python Implementation of Codon Adaptation Index. J. Open Source Softw. 3:905.

Lee SJ, Lee S-R. 2022. Complete mitochondrial genome of the abalone shell-boring Polydora hoplura (Polychaeta, Spionidae). Mitochondrial DNA B Resour. 7:438–439.

Manchenko GP, Radashevsky VI. 1993. Genetic differences between two sibling species of the Polydora ciliata complex (Polychaeta: Spionidae). Biochem. Syst. Ecol. 21:543–548.

Mangkalaphiban K et al. 2021. Transcriptome-wide investigation of stop codon readthrough in Saccharomyces cerevisiae. PLoS Genet. 17:e1009538.

Martijn J, Vosseberg J, Guy L, Offre P, Ettema TJG. 2018. Deep mitochondrial origin outside the sampled alphaproteobacteria. Nature. 557:101–105.

Meng G, Li Y, Yang C, Liu S. 2019. MitoZ: a toolkit for animal mitochondrial genome assembly, annotation and visualization. Nucleic Acids Res. 47:e63.

Milani L, Ghiselli F, Guerra D, Breton S, Passamonti M. 2013. A comparative analysis of mitochondrial ORFans: new clues on their origin and role in species with doubly uniparental inheritance of mitochondria. Genome Biol. Evol. 5:1408–1434.

Mitchell A, Guerra D, Stewart D, Breton S. 2016. In silico analyses of mitochondrial ORFans in freshwater mussels (Bivalvia: Unionoida) provide a framework for future studies of their origin and function. BMC Genomics. 17:597.

Mustaquim J. 1986. Morphological variation in Polydora ciliata complex (Polychaeta:Annelida). Zool. J. Linn. Soc. 86:75–88.

Nei M, Gojobori T. 1986. Simple methods for estimating the numbers of synonymous and nonsynonymous nucleotide substitutions. Mol. Biol. Evol. 3:418–426.

Pagès H, Aboyoun P, R. G, DebRoy S. 2017. Biostrings. Bioconductor doi:10.18129/B9.BIOC.BIOSTRINGS.

Pons J-L, Labesse G. 2009. @TOME-2: a new pipeline for comparative modeling of protein-ligand complexes. Nucleic Acids Res. 37:W485–91.

Pont-Kingdon G et al. 1998. Mitochondrial DNA of the coral Sarcophyton glaucum contains a gene for a homologue of bacterial MutS: a possible case of gene transfer from the nucleus to the mitochondrion. J. Mol. Evol. 46:419–431.

Pont-Kingdon GA et al. 1995. A coral mitochondrial mutS gene. Nature. 375:109–111.

Potter SC et al. 2018. HMMER web server: 2018 update. Nucleic Acids Res. 46:W200–W204.

Raboin MJ, Timko AF, Howe DK, Félix M-A, Denver DR. 2010. Evolution of Caenorhabditis mitochondrial genome pseudogenes and Caenorhabditis briggsae natural isolates. Mol. Biol.Evol. 27:1087–1096.

Radashevsky VI, Migotto AE. 2017. First report of the polychaete Polydora hoplura (Annelida: Spionidae) from North and South America and Asian Pacific. Mar. Biodivers. 47:859–868.

Radashevsky VI, Pankova VV. 2013. Shell-boring versus tube-dwelling: is the mode of life fixed or flexible? Two cases in spionid polychaetes (Annelida, Spionidae). Mar. Biol. 160:1619–1624.

Radashevsky VI, Pankova VV. 2006. The morphology of two sibling sympatric Polydora species (Polychaeta: Spionidae) from the Sea of Japan. J. Mar. Biol. Assoc. U. K. 86:245–252.

Ranwez V, Harispe S, Delsuc F, Douzery EJP. 2011. MACSE: Multiple Alignment of Coding SEquences accounting for frameshifts and stop codons. PLoS One. 6:e22594.

San Mauro D, Gower DJ, Zardoya R, Wilkinson M. 2006. A hotspot of gene order rearrangement by tandem duplication and random loss in the vertebrate mitochondrial genome. Mol. Biol. Evol. 23:227–234.

Sato-Okoshi W, Abe H, Nishitani G, Simon CA. 2017. And then there was one: Polydora uncinata and Polydora hoplura (Annelida: Spionidae), the problematic polydorid pest species represent a single species. J. Mar. Biol. Assoc. U. K. 97:1675–1684.

Schultz DT et al. 2020. Conserved novel ORFs in the mitochondrial genome of the ctenophore Beroe forskalii. PeerJ. 8:e8356.

Soldatov RA, Vinogradova SV, Mironov AA. 2014. RNASurface: fast and accurate detection of locally optimal potentially structured RNA segments. Bioinformatics. 30:457–463.

Sveshnikov VA. 1958. New species of Polychaetes (Annelida) for the White Sea. (In Russian). Zoologichesky Zhurnal. 37:20–26.

Tamura K, Stecher G, Kumar S. 2021. MEGA11: Molecular Evolutionary Genetics Analysis Version 11. Mol. Biol. Evol. 38:3022–3027.

Tassé M et al. 2022. The longest mitochondrial protein in metazoans is encoded by the male-transmitted mitogenome of the bivalve Scrobicularia plana. Biol. Lett. 18:20220122.

Waterhouse AM, Procter JB, Martin DMA, Clamp M, Barton GJ. 2009. Jalview Version 2--a multiple sequence alignment editor and analysis workbench. Bioinformatics. 25:1189–1191.

Weigert A et al. 2016. Evolution of mitochondrial gene order in Annelida. Mol. Phylogenet.Evol. 94:196–206.

Wickham H. 2016. ggplot2. Springer International Publishing.

Wickham H et al. 2019. Welcome to the tidyverse. J. Open Source Softw. 4:1686.

Wickham H, François R, Henry L, Müller K. 2022. dplyr: A Grammar of Data Manipulation.dplyr: A Grammar of Data Manipulation. https://dplyr.tidyverse.org/authors.html (Accessed November 30, 2022).

Ye L, Yao T, Lu J, Jiang J, Bai C. 2021. Mitochondrial genomes of two Polydora (Spionidae)species provide further evidence that mitochondrial architecture in the Sedentaria (Annelida)is not conserved. Sci. Rep. 11:13552.

Zheng W et al. 2021. Folding non-homologous proteins by coupling deep-learning contact maps with I-TASSER assembly simulations. Cell Rep Methods. 1. doi:10.1016/j.crmeth.2021.100014.

Zimmermann L et al. 2018. A Completely Reimplemented MPI Bioinformatics Toolkit with a New HHpred Server at its Core. J. Mol. Biol. 430:2237–2243.

