## Supplementary materials for "ORFanes in mitochondrial genomes of marine polychaete *Polydora*"

**Table S1.** Long range PCR primers. These primers were used to enrich mitochondrial DNAs, PCR products were sequenced by Illumina, reads were mapped to the assembly and used to verify it.

| Name | Primer | Tm |
|---|---|---|
| F1 | TTTTGAACGCCATGAGGGGG | 60 |
| R1 | GGTTCATCCAGTCCCTGCTC | 60 |
| F2 | TACTCTCTCCCTCTTTCGCCA | 59 |
| R2 | AGGTGCAAGCTAGATGTTCTATTT | 59 |
| F3 | CATCTGGAGCTCCGTCAACA | 60 |
| R3 | CCTGCTCACCCTCATTGTGT | 60 |
| F4 | CGACGCAGGACTTCCTTGTT | 60 |
| R4 | ATCTTCAGTGTGGCGCTCTT | 60 |

**Table S2.** Primers flanking unannotated regions. These primers were used to confirm sequences of ORFanes and CR using Sanger sequencing.

| Name | Primer | Tm | Region |
|---|---|---|---|
| F1 | TCCTCAACAGICICCAAICCA | 57 | 1245-1268 |
| R1 | ATAGGAGCTGAAGGGGACAT | 57 | 1991-1972 |
| F2 | CGCCCAGGTACAGTCTTTGT | 59 | 8716-8735 |
| R2 | CTCAATGTTGGGGCATGAC | 59 | 9474-9455 |
| F3 | CCTTTGACATCCGAAGTATAG GTA | 58 | 11648-11671 |
| R3 | ATAAGCGTTTACCCGAGCAC | 58 | 12906-12887 |
| F4 | TTCCCTATCCCCTTAGCACT | 58 | 13856-13857 |
| R4 | GCAAGGCCTAGGAGAGACC | 58 | 14799-14780 |

**Table S3.** Atome3 and I-tasser top 10 hits (for Atome3 score threshold of 40.00 also applied, for I-tasser duplicate hits for one protein are excluded) of the mitochondrial *Polydora* and *Boccardiella* species ORFans.

| ORFan1 | ORFan2 | ORFan3 |
|---|---|---|
| Atome3 (PDB id, Classification, score, identity) | | |
| ***P. cf. ciliata***<br>4LXU, transferase, 86.04, 18%<br>3ZPJ, unknown function, 71.06, 15<br>1GZ6, dehydrogenase, 67.09, 11%<br>6EU6, membrane protein, 43.64, 30%<br>2OKQ, unknown function, 42.09, 14%,<br>7E2I, lipid transport, 40.73, 30%<br>7E2G, lipid transport, 40.59, 30%<br><br>***P. websteri***<br>3IBT, oxidoreductase, 72.46, 18%<br>3OQ4, cell cycle, 45.68, 9%<br>4TT0, hydrolase, 44.59, 10%<br>2AHQ, transcription, 40.19, 16%,<br>7M8W, membrane protein, 39.94, 33%<br><br>***P. brevipalpa***<br>5LRT, hydrolase, 84.97, 18%<br>5I3E, hydrolase, 68.57, 16%<br>5FGU, Metal binding, DNA binding protein, 59.54, 17%<br>3FWK, transferase, 48.62, 13%<br><br>***P. hoplura***<br>5GAP, transcription, 75.96, 17%<br>4RDQ, transport protein, 73.68, 16%<br>4KYI, protein binding/transport protein, 68.26, 15%<br>1PV6, transport protein, 62.45, 9%<br>5FT3, transferase, 53.58, 15%<br><br>***B. hamata***<br>*5T9J,* hydrolase, 77.36, 16%<br>3ALX, viral protein/membrane protein, 75.83, 23%<br>5AHR, hydrolase, 71.38, 22%<br>4BWZ, transport protein, 65.06, 14%<br>5IOJ, hydrolase, 62.28, 16%<br>4IIK, hydrolase, 61.81, 13%<br>3GIA, transport protein, 59.87, 10%<br>5CA8, hydrolase, 59.73, 16%<br>4L4W, protein transport, 58.55, 17%<br>5TCQ, membrane protein, 56.29, 18% | ***P. cf. ciliata***<br>*2KCD,* unknown function, 77.50, 21%<br>2MN4, de novo protein, 66.92, 20%<br>5JK2, cell adhesion, 64.44, 16%<br>3E0Z, unknown function, 61.57, 20%<br>4DQJ, hydrolase, 60.02, 13%<br>3A98, signaling protein, 59.51, 16%<br>2WCR, immune system, 58.11, 16%<br>4P02, transferase, 50.63, 26%<br>2JKG, protein binding, 49.48, 12%<br>6V93, DNA binding protein, 49.42, 25%<br><br>***P. websteri***<br>3FRR, protein binding, 72.44, 11%<br>4NQI, signaling protein, 64.86, 185<br>2EE3, signaling protein, 64.60, 15%<br>4P02, transferase, 61.76, 31%<br>2I9S, chaperone, 59.82, 17%<br>5EJ1, metal binding protein, 58.13, 30%<br>5IFG, hydrolase/antitoxin, 57.77, 13%<br>5E6G, de novo protein, 56.29, 11%<br>6UEB, viral protein, score 53.10, 25%<br>2HEQ, unknown function, 50.24, 19%<br><br>***P. brevipalpa***<br>*3A98,* signaling protein, 71.18, 20%<br>2MN4, de novo protein, 60.39, 22%<br>7BJK, oxidoreductase, 59.34, 27%<br>2DDZ, unknown function, 51.35, 11%<br>2FM9, cell invasion, 49.77, 11%<br>4P02, transferase, 43.47, 29%<br>5EJ1, metal binding protein, 42.24, 29%<br>1VI7, unknown function, 41.78, 15%<br>4P1Z, RNA binding protein, 40.57, 10%<br><br>***P. hoplura***<br>*5M11,* transport protein, 66.24, 21%<br>3A98, signaling protein, 61.49, 14%<br>4R6I, transcription, 60.98, 19%<br>4ZOX, chaperone, 60.33, 13%<br>5IFG, hydrolase/antitoxin, 57.83, 17%<br>2EE3, signaling protein, 55.40, 17%<br>2KCD, unknown function, 51.55, 18%<br>1JHU, transferase, 50.97, 6%<br>6WLZ, membrane protein, 40.35, 42%<br><br>***B. hamata***<br>1UG7, structural genomics/unknown function, 86.09, 19%<br>6BWI, membrane protein, 72.75, 30%<br>6BQV, membrane protein, 62.20, 30%<br>6BCO, transport protein, 62.03, 30%<br>6BQR, 61.23, 30%<br>5WP6, membrane protein, 59.71, 30%<br>2EDB, apoptosis, 58.28, 20% | ***P. cf. ciliata***<br>*4LEU,* RNA binding protein, 78.89, 13%<br>5JJO, immune system, 78.23, 19%<br>4S2R, hydrolase, 70.71, 24%<br>3N0K, hydrolase inhibitor, 70.52, 14%<br>2B6C, unknown function, 66.91, 12%<br>2C0N, viral protein/transferase, 65.40, 17%<br>2VLI, transferase, 62.85, 10%<br>4WIA, ATP-binding protein, 62.43, 11%<br>2QEC, transferase, 61.64, 11%<br>1CA4, TNF signaling, 58.32, 11%<br><br>***P. websteri***<br>*1WCK,* signaling protein,, 70.75, 15%<br>5UGW, transferase, 59.83, 15%<br>2VLI, transferase, 56.06, 9%<br>1ZWT, cell adhesion, 50.49, 20%<br>1J3G, hydrolase, 49.91, 13%<br>2EDO, cell adhesion, 49.75, 14%<br>2E33, lactase/hydrolase, 49.48, 25%<br>1UMI, ligase, 48.42, 25%<br>2E31, ligase, 48.12, 25%<br>1UMH, ligase, 47.95, 25%<br>2RJ2, ligase, 47.58, 25%<br><br>***P. brevipalpa***<br>2OH5, structure protein/RNA binding protein, 68.10, 18%<br>3G8Q, RNA binding protein, 67.52, 18%<br>2AN1, transferase, 61.52, 11%<br>5LQ6, immunosuppressant, 58.64, 14%<br>3N0K, hydrolase inhibitor, hydrolase, 14%<br>2WWX, protein transport, 54.03, 18%<br>4G6V, toxin, 53.12, 15%<br>4NQI, signaling protein, 52.97, 12%<br>3HG9, unknown function, 50.09, 17%<br>4FPR, protein binding, 49.00, 20%<br><br>***P. hoplura***<br>3G8Q, RNA binding protein, 71.71, 16%<br>3E9C, hydrolase, 66.33, 15%<br>3DCM, transferase, 64.91, 14%<br>3WG9, transcription, 61.86, 16%<br>5ANB, translation, 57.21, 18%<br>2AMJ, oxidoreductase, 54.93, 10%<br>2FO1, gene regulation/signaling protein, 54.28, 18%<br>1WBL, lectin, 52.85, 15% |

| | | |
|---|---|---|
| | 1BCF, iron storage and electron transport, 57.79, 9%<br>3WX4, viral protein, 56.92, 12%<br>2ND4, hydrolase receptor, 54.61, 11% | 2C0N, viral protein/transferase, 47.76, 19%<br>2OST, hydrolase, 46.40, 15%<br><br>***B. hamata***<br>4QGN, oxidoreductase, 73.33, 13%<br>5A9H, transport protein, 72.68, 13%<br>1CA4, TNF signaling, 71.32, 11%<br>5FT0, hydrolase inhibitor, 68.88, 18%<br>5B5Z, metal binding protein, 65.53, 14%<br>3KEA,viral protein, 64.49, 19%<br>5ARM, copper-binding protein, 63.63, 13%<br>4PHR, transferase, 61.79, 18% |
| I-tasser (PDB id, Classification, identity1, identity2, coverage) | | |
| ***P. cf. ciliata***<br>3H1I, oxidoreductase, 0.18, 0.18, 0.82<br>2PFF, transferase, 0.40, 0.27, 0.06<br>6FVB, nuclear protein, 0.16, 0.30, 0.78<br>7DBG, transport protein, 0.14, 0.19, 0.92<br>3ZKV, transport protein, 0.12, 0.28, 0.86<br>7MEX, transferase, 0.13, 0.27, 0.95<br>2XWU, ligase/nuclear protein, 0.13, 0.28, 0.91<br>7WSS, hydrolase, 0.13, 0.21, 0.97<br>5NVR, structural protein, 0.16, 0.29, 0.93<br>7QJ0, cytosolic protein, 0.12, 0.22, 0.94<br><br>***P. websteri***<br>4RY2, transport protein/hydrolase, 0.14, 0.20, 0.83<br>2PFF, transferase, 0.29, 0.28, 0.81<br>6FVB, nuclear protein, 0.13, 0.30, 0.96<br>7DBG, transport protein, 0.12, 0.20, 0.91<br>3QF4, transport protein, 0.12, 0.17, 0.81<br>3JAC, metal transport, 0.19, 0.26, 0.91<br>6N1Z, transport protein, 0.11, 0.26, 0.86<br><br>***P. brevipalpa***<br>4RY2, transport protein/hydrolase, 0.10, 0.22, 0.82<br>5T8V, cell cycle, 0.15, 0.23, 0.88<br>7DBG, transport protein, 0.16, 0.19, 0.89<br>6KG7, membrane protein, 0.16, 0.17, 0.88<br>5WYL, ribosomal protein/nuclear protein, 0.16, 0.20, 0.71<br>7WKK, structural protein, 0.15, 0.22, 0.84<br>5DLQ, protein transport, 0.16, 0.30, 0.88<br>7OCI, membrane protein, 0.15, 0.25, 0.96<br>5U1T, hydrolase, 0.18, 0.32, 0.79<br>7QE7, cell cycle, 0.17, 0.21, 0.94<br><br>***P. hoplura***<br>*6WW2,* membrane protein, 0.12, 0.24, 0.88<br>2PFF, transferase, 0.27, 0.25, 0.13<br>5UFL, signaling protein, 0.17, 0.25, 0.88<br>7DBG, transport protein, 0.14, 0.18, 0.92<br>5NL2, membrane protein, 0.14, 0.20, 0.86<br>6FVB, nuclear protein, 0.13, 0.35, 0.91 | ***P. cf. ciliata***<br>4FGV, transport protein, 0.15, 0.16, 0.75<br>7WKK, structural protein, 0.11, 0.50, 0.99<br>1HS6, hydrolase, 0.18, 0.19, 0.92<br>3JAC, metal transport, 0.18, 0.31, 0.41<br>5H3O, transport protein, 0.13, 0.22, 0.95<br>5NL2, membrane protein, 0.11, 0.17, 0.98<br>4V4L, apoptosis, 0.25, 0.35, 0.98<br>5UFK, protein binding, 0.15, 0.25, 0.86<br>7ZCV, DNA binding protein, 0.12, 0.30, 0.99<br><br>***P. websteri***<br>6W6X, de novo protein, 0.14, 0.17, 0.77<br>7M68, membrane protein, 0.09, 0.42, 0.98<br>2MN2, antitoxin, 0.18, 0.18, 0.87<br>4HG6, transferase, 0.32, 0.23, 0.44<br>7LP9, membrane protein, 0.08, 0.17, 0.96<br>5NL2, membrane protein, 0.08, 0.22, 0.1<br>5DN6, hydrolase, 0.15, 0.12, 0.45<br>5WZJ, RNA binding protein/RNA, 0.14, 0.30, 0.96<br>7ZCV, DNA binding protein, 0.15, 0.25, 0.96<br><br>***P. brevipalpa***<br>4MU6, unknown function, 0.16, 0.19, 0.88<br>7W7G, membrane protein, 0.09, 0.24, 0.90<br>4ZQB, oxidoreductase, 0.16, 0.26, 0.90<br>5MZ4, viral protein, 0.53, 0.26, 0.12<br>6O02, transferase/protein binding 0.25, 0.18, 0.86<br>3JBR, membrane protein, 0.08, 0.13, 1.0<br>2PFF, transferase, 0.25, 0.32, 0.91<br>5UFK, protein binding, 0.21, 0.28, 0.86<br>7ZCV, DNA binding protein, 0.15, 0.18, 0.94 | ***P. cf. ciliata***<br>4N5Q, protein binding, 0.11, 0.18, 0.72<br>7SZX, viral protein, 0.10, 0.17, 0.85<br>6Q8J, splicing, 0.14, 0.19, 0.90<br>4V4L, apoptosis, 0.37, 0.38, 0.15<br>4ZGN, cell cycle, 0.17, 0.17, 0.72<br>4DR0, oxidoreductase, 0.18, 0.24, 1.0<br>3JAC, metal transport, 0.18, 0.27, 0.93<br>6DJY, virus, 0.11, 0.37, 0.78<br>7VOI, hydrolase, 0.22, 0.25, 0.89<br><br>***P. websteri***<br>2J5T, transferase, 0.14, 0.16, 0.88<br>7NQD, transferase, 0.06, 0.19, 0.89<br>4C47, cell adhesion, 0.20, 0.19, 0.94<br>1D3Y, isomerase, 0.19, 0.14, 0.26<br>6LF6, transferase, 0.24, 0.17, 0.68<br>6ZP9, viral protein, 0.09, 0.17, 0.96<br>4V4L, apoptosis, 0.27, 0.16, 0.76<br>5UN8, hydrolase, 0.18, 0.27, 0.85<br>7P6G, hydrolase, 0.15, 0.23, 0.92<br><br>***P. brevipalpa***<br>5YO8, lyase, 0.15, 0.12, 0.*67*<br>*7BZX, transferase, 0.13, 0.28, 0.94*<br>*5ZMO, DNA binding protein/DNA, 0.13, 0.20, 0.82*<br>2PFF, transferase, 0.20, 0.36, 0.16<br>7KB3, signaling protein, 0.20, 0.20, 0.97<br>6W2V, biosynthetic protein, 0.11, 0.16, 0.98<br>2R7G, transcription repressor/cell cycle, 0.19, 0.25, 0.70<br>7VOI, hydrolase, 0.20, 0.23, 0.94<br><br>***P. hoplura***<br>2PFF, transferase, 0.31, 0.32, 0.85<br>4V4L, apoptosis, 0.25, 0.18, 0.71<br>2LF6, signaling protein,0.21, 0.14, 0.43<br>7FIV, protein binding, 0.12, 0.16, 0.78<br>1H6U, cell adhesion, 0.17, 0.20, 0.96<br>3CJH, protein transport, 0.23, 0.07, 0.14 |

| | | |
|---|---|---|
| 7WKK, structural protein, 0.15, 0.23, 0.95<br>7P5C, lipid transport, 0.10, 0.21, 0.81<br>5OQQ, cell cycle, 0.18, 0.27, 0.80<br><br>***B. hamata***<br>*3JAC, metal transport, 0.21, 0.25, 0.88*<br>*2XWU,* ligase/nuclear protein, 0.14, 0.29, 0.58<br>7OCI, membrane protein, 0.14, 0.27, 0.99<br>4V4L, apoptosis, 0.24, 0.12, 0.42<br>4HAT, protein transport/antibiotic, 0.22, 0.33, 0.57<br>7W7G, membrane protein, 0.18, 0.23, 0.92<br>1W27, lyase, 0.15, 0.27, 0.85<br>7ESI, hydrolase, 0.14, 0.21, 0.95 | ***P. hoplura***<br>1ZKR, allergen, 0.14, 0.17, 0.82<br>7M68, membrane protein, 0.12, 0.42, 1.0<br>4EJT, transcription regulator/RNA<br>2AFF, cell cycle, 0.53, 0.08, 0.11<br>4XMN, protein transport, 0.17, 0.19, 0.91<br>6XR1, de novo protein, 0.07, 0.19. 1.0<br>3JAC, metal transport, 0.15, 0.30, 0.98<br>6R7O, gene regulation, 0.17, 0.26, 0.96<br>7VOI, hydrolase, 0.19, 0.26, 0.95<br><br>***B. hamata***<br>1T33, transcription, 0.13, 0.15, 0.79<br>7N0A, cytokine/immune system, 0.14, 0.19, 0.99<br>6TV5, structural protein, 0.15, 0.18, 0.94<br>6TGB, signaling protein, 0.17, 0.12, 0.50<br>2MX8, structural protein, 0.09, 0.17, 0.90<br>6BQ1, transferase/signaling protein, 0.20, 0.42, 0.98<br>6IGX, cell cycle, 0.13, 0.38, 1.0<br>7U7N, cytokine, 0.15, 0.16, 0.96 | 6HXZ, virus like particle, 0.24, 0.17, 0.81<br>6W2V, biosynthetic protein, 0.07, 0.16, 0.97<br>3JAC, metal transport, 0.17, 0.33, 0.80<br><br>***B. hamata***<br>2PFF, transferase, 0.33, 0.31, 0.11<br>4V4L, apoptosis, 0.24, 0.31, 0.89<br>1HRK, lyase, 0.16, 0.16, 0.69<br>7NHA, viral protein, 0.16, 0.21, 0.87<br>4R7Q, signaling protein, 0.20, 0.22, 0.96 |

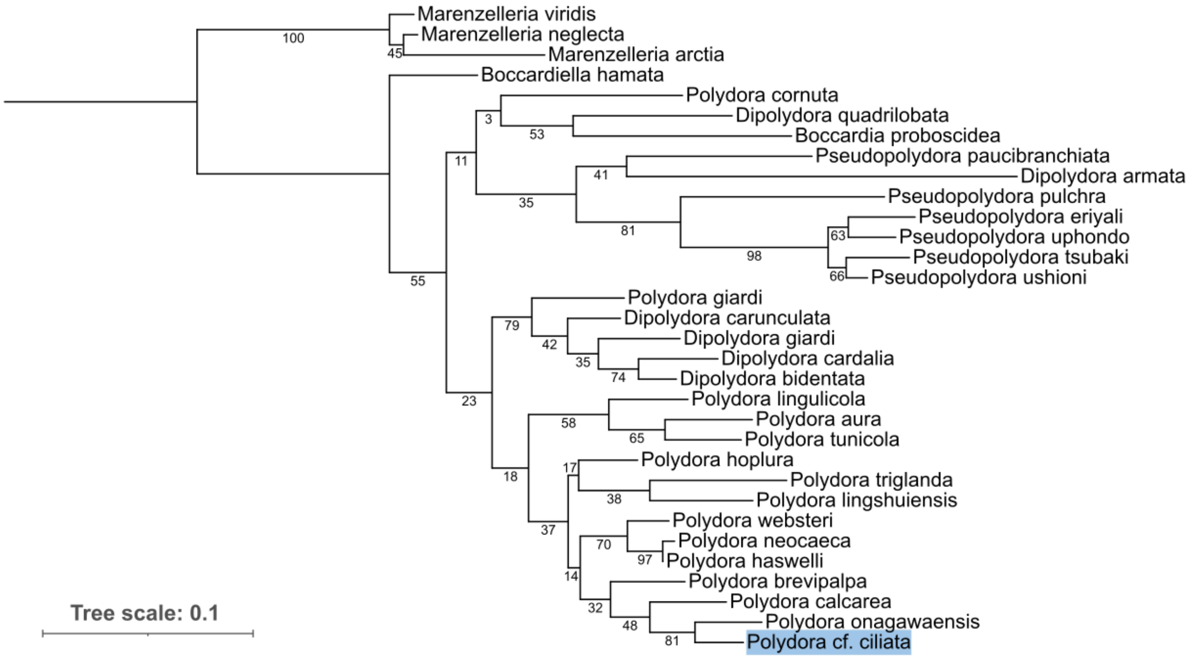


**Figure S1.** Phylogenetic tree of 16S sequences of *Polydora cf. ciliate* and related Annelida species constructed using the Maximum Likelihood Algorithm with General Time Reversible model integrated in MEGA11 software with default parameters and 100 bootstrap replicas.

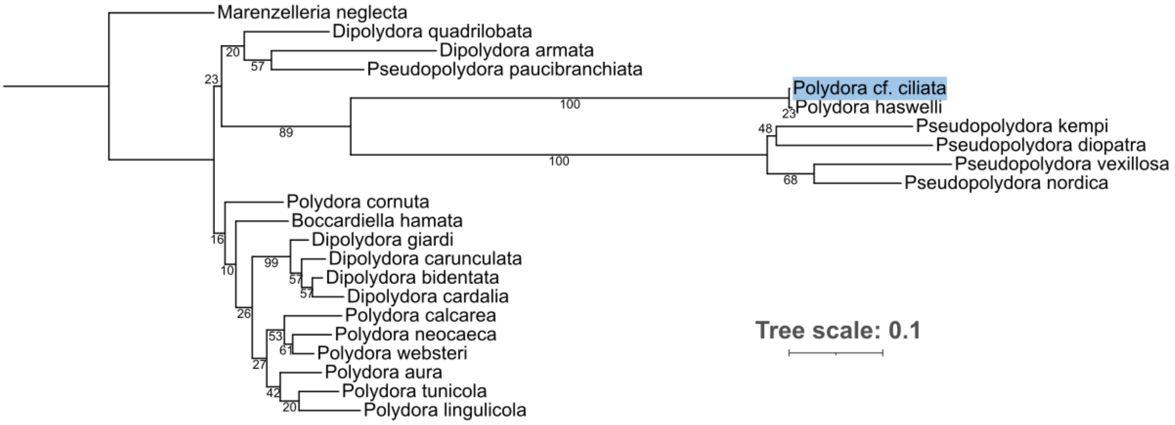


**Figure S2.** Phylogenetic tree of 18S sequences of *Polydora cf. ciliate* and related Annelida species constructed using the Maximum Likelihood Algorithm with a General Time Reversible model integrated in MEGA11 software with default parameters and 100 bootstrap replicas.

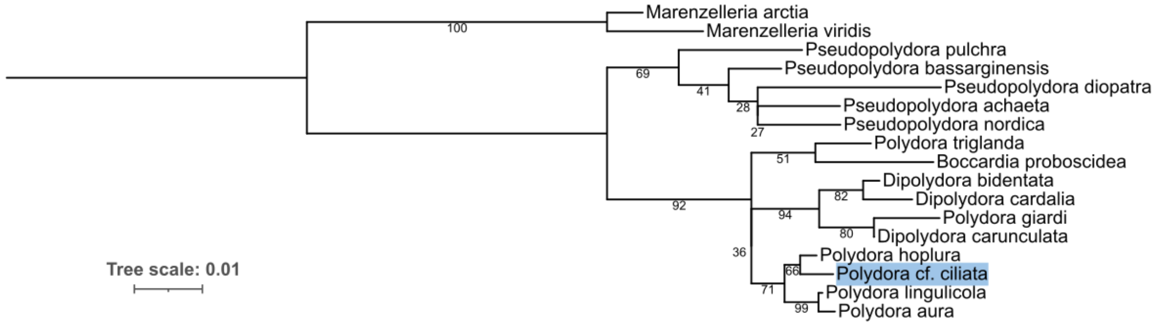


**Figure S3.** Phylogenetic tree of 28S sequences of *Polydora cf. ciliate* and related Annelida species constructed using the Maximum Likelihood Algorithm with a General Time Reversible model integrated in MEGA11 software with default parameters and 100 bootstrap replicas

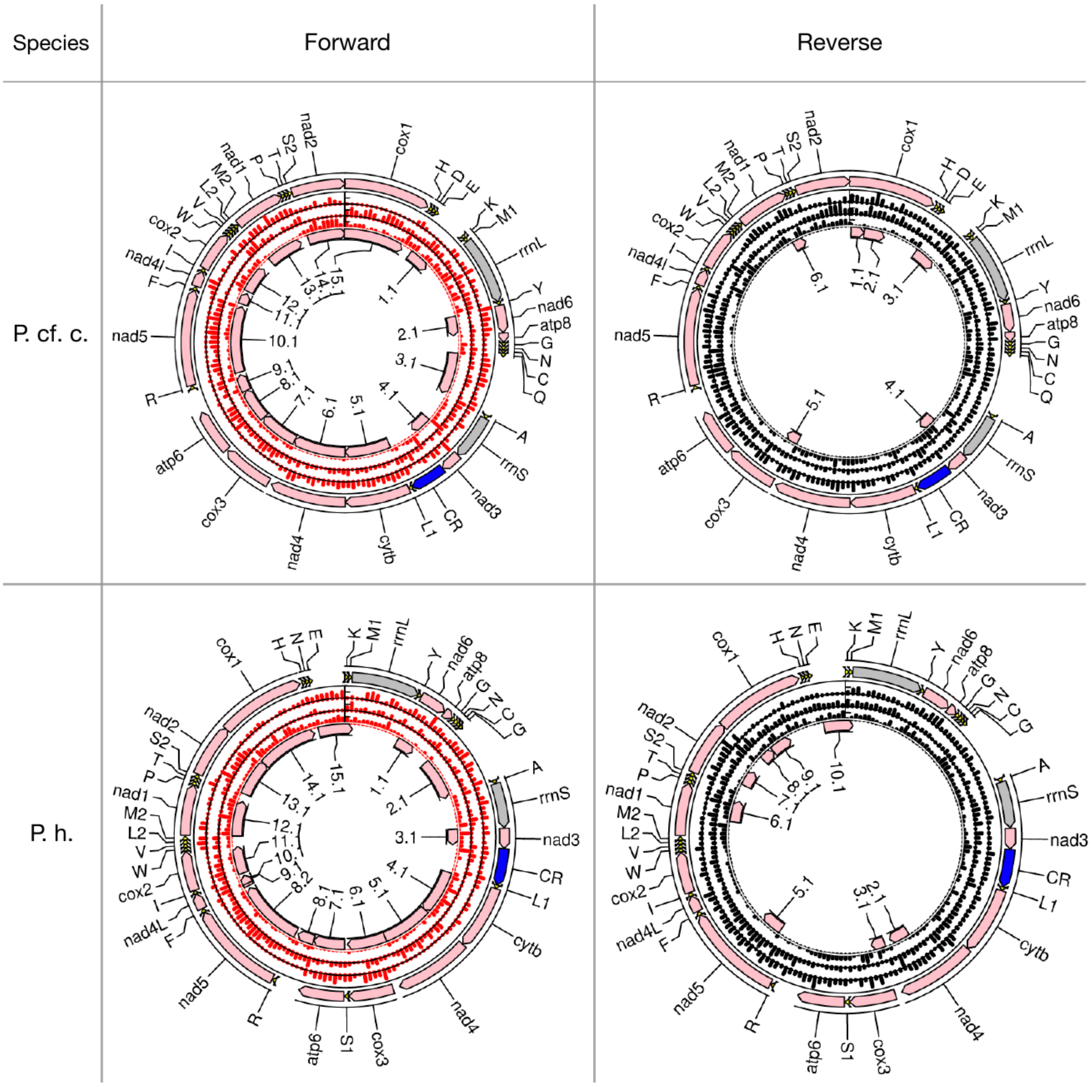

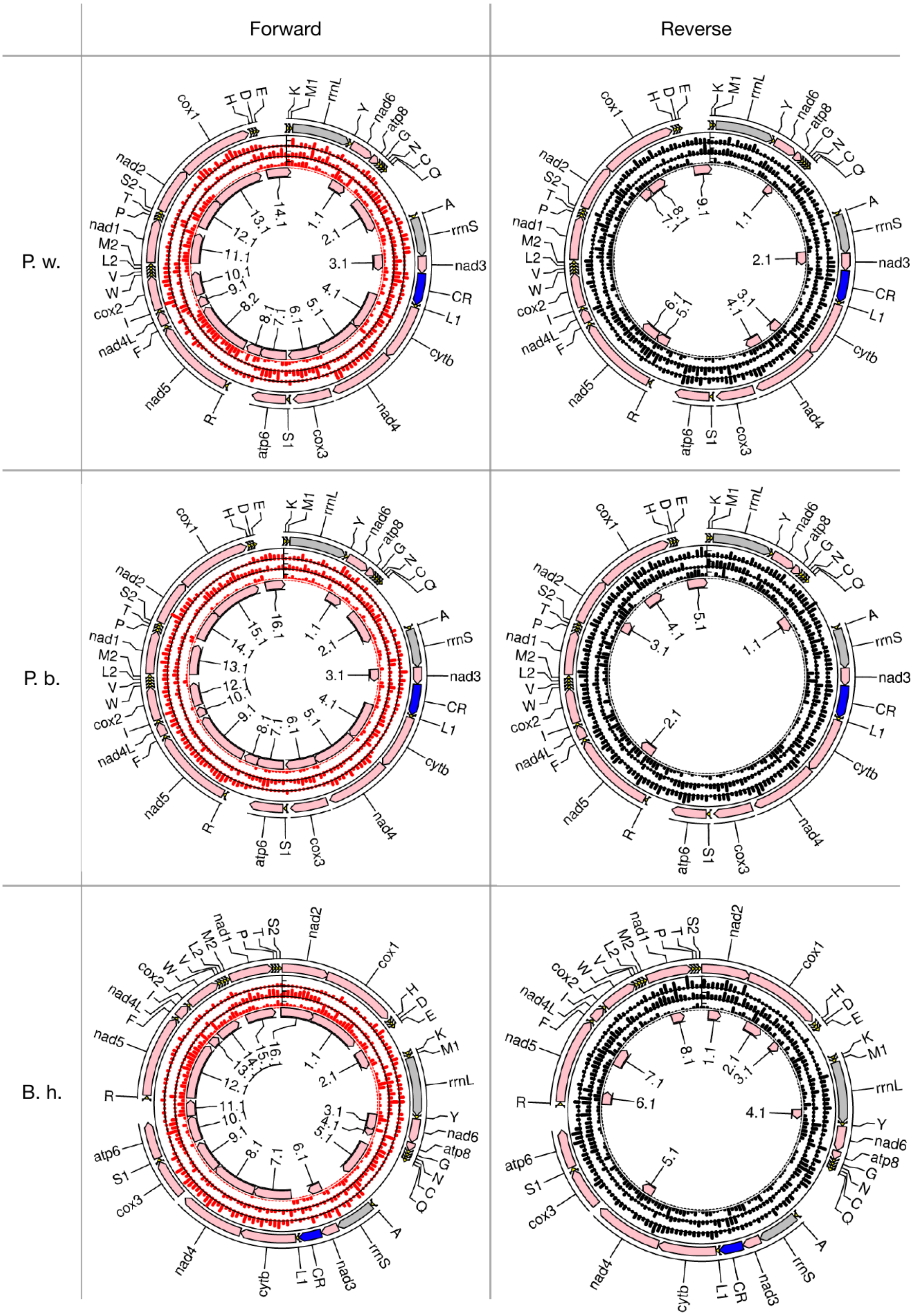

Forward
Reverse
P. w.
P. b.
B. h.

**Figure S4. Maps of all analysed mitogenomes with visualised stop codons in all six frames.** Outer circle shows annotation for a particular genome, three mid-circles show the amount of stop codons per bin for each frame, where red indicates stop codons in forward strand, black in reverse strand. Inner circle shows potential protein-coding regions after tRNA-punctuation of long regions lacking stop-codons (see methods). Species transcript: *P. cf. c. – Polydora cf. ciliata, P. h. – Polydora hoplura, P. w. – Polydora websteri, P. b. – Polydora brevpalpa, B. h. – Boccardiella hamata.*

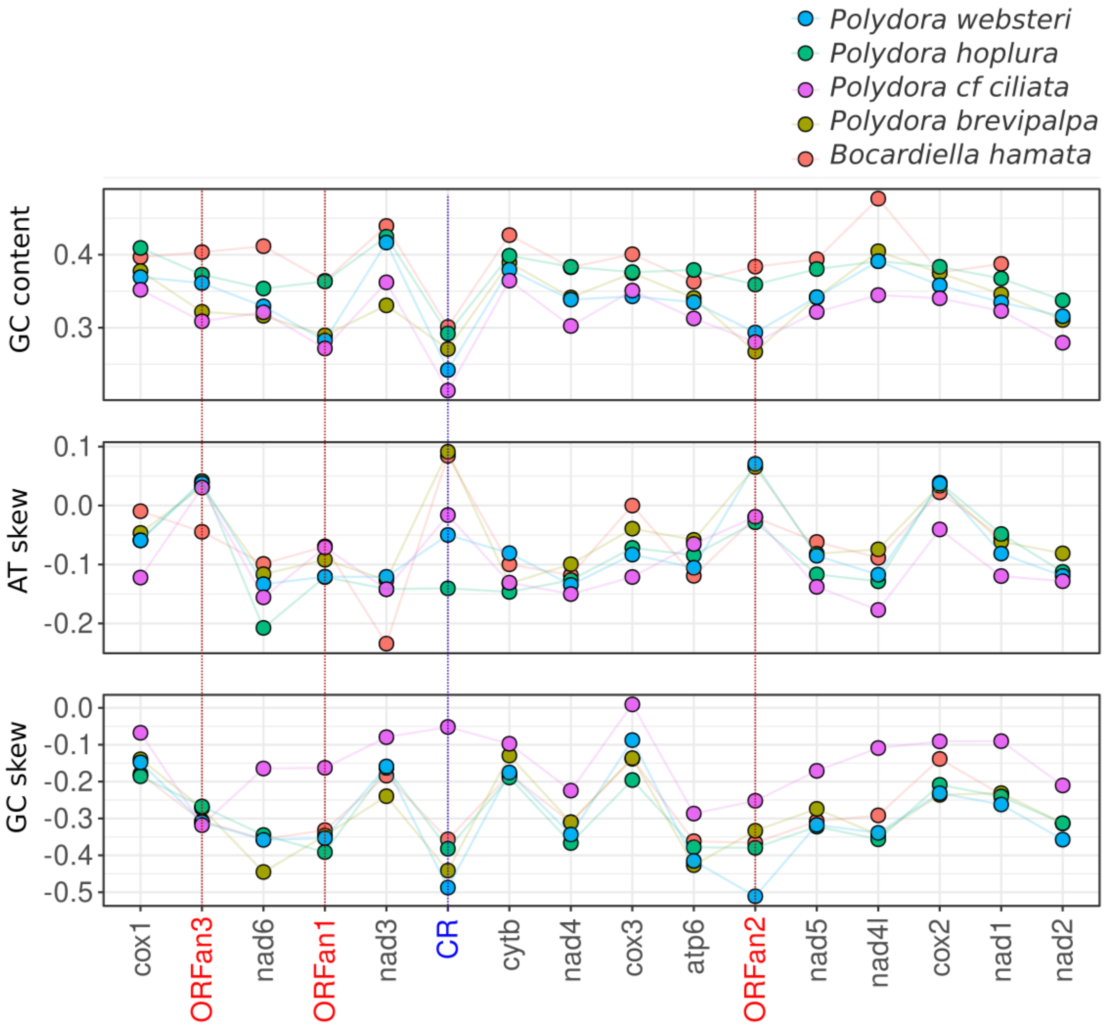

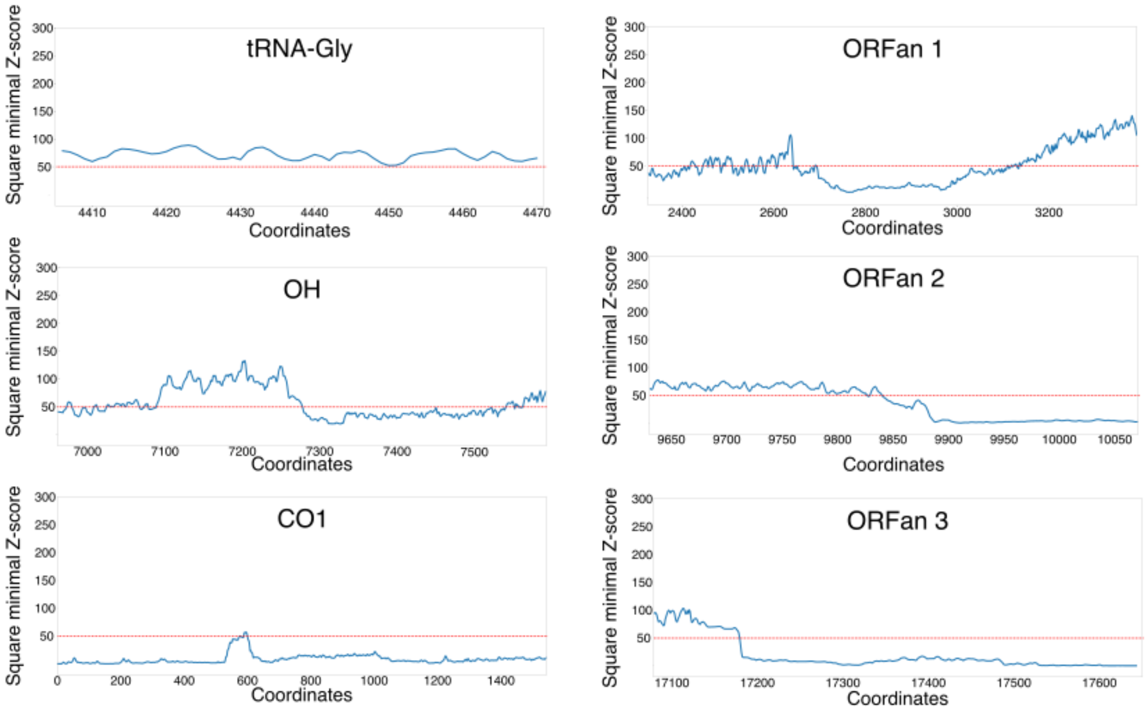


**Figure S5. A.** Analysis of the nucleotide content of *Polydora* and Bocardiella mitochondrial genes and putative control region (CR) (GC-content, A-T and G-C skew) **B.** Analysis of the nucleotide sequence for its ability to form secondary structures in a single-stranded state. We analysed three ORFan genes, tRNA-Gly, CO1 and a possible control region. The ability of a sequence to form a stable secondary structure was measured by computing Z-scores; higher values of square minimum Z-score reflect higher significance of secondary structure in the region. The possible presence of regulatory secondary structures was addressed using RNASurface server (Soldatov, Vinogradova, and Mironov 2014).

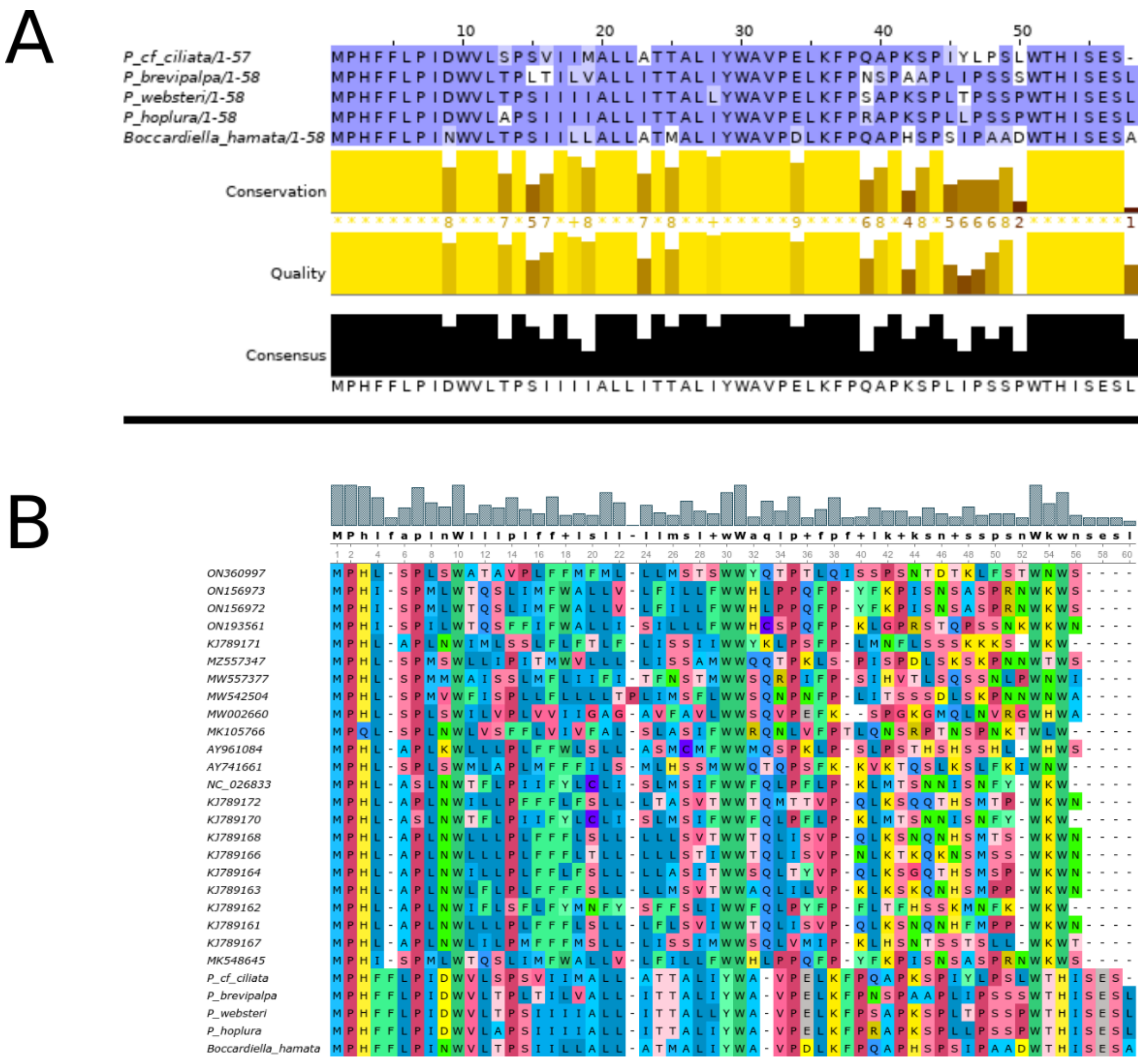


**Figure S6.** Multiple alignments of ATP8 genes. Given the highly conservative nature of mitogenome architecture in *Polydora* genus and *Bocardiella hamata* we searched for atp8 right after nad6 in the same position where the gene was located in species with annotated atp8. We performed multiple alignment of atp8 genes from *Polydora* and *Bocardiella* species [Muscle with defaults parameters, JalView] (A). The sequences proved to be nearly identical, thus we concluded that these ORFs are indeed atp8 sequences not found by automatic annotation. We conducted multiple alignment of all known sedentarian atp8 sequences [using Muscle algorithm with defaults parameters] (B). The alignment revealed substitution of the conserved part of the protein: the first four residues consensus appears to be MPHL instead of MPQL. Although we see the same three first a.a. residues in *Polydora* genus, its atp8 exhibits exceptional differences from other sedentarian species.

### Supplementary text 1. Description of remote homology search algorithms outputs.

In order to detect more remote homologies we also performed HMM vs sequence search using hmmsearch against available databases (see methods). As a result of the search, no significant hits (with the cutoff value of 0.01 as suggested by the program) were found in any of the databases for both the first and the second ORFan proteins. For the third ORFan protein, hmmsearch found one significant hit (e-value=0.0028) in the ‘reference proteomes' database and two significant hits in the UniProtKB (e-values=0.0009 and 0.0083).

All hits were “uncharacterised proteins” either from *Tenericutes bacterium* (recommended name: CARDB domain-containing protein) or *Cinara cedri* (recommended name: RNA-directed DNA polymerase). HMM vs HMM search performed by HHpred revealed several possible homologies, yet with a probability not exceeding 70%. As suggested by HHpred we considered hits with > 50% and hits which are among top three hits with > 30% probability. Given the suggested cutoff three hits for ORFan1 were found: RNA-binding protein 42 from *Homo sapiens*, Putative gene 60 protein from *Bacillus subtilis* and Uncharacterized protein yaiA from *Escherichia coli*. These hits hardly indicate the function of the protein studied. For ORFan2 21 hits were found while top seven hits with similar probability (approximately 60%) were proteins connected with ABC transporters from different bacteria. Among other hits there also were several transport and membrane proteins. These findings suggest that ORFan2 may be connected with membrane transport. No hits were found for ORFan3.

Tertiary structure prediction and search for structure similarity for three ORFanes in five species was performed using @tome. For ORFan1 there were proteins connected with the hydrolase family in all species studied. However, the level of similarity does not exceed 40% and the significance score does not exceed 30 in none of the cases. Some hits are connected with transmembrane proteins associated with transport, especially in *Polydora cf. ciliata*. For ORFan2 transferases or parts of transferases were found in all species. Although these findings were insignificant, these results are consistent with the results of HHpred which pointed on ABC transporters. For ORFan3 there were hydrolases, transferases and dna-binding proteins among hits in all species, however percentage identity and score were substantially low. Interestingly, @tome found a signal peptide in ORFan3 of Polydora brevipalpa which is consistent with TOPCONs results even though the prediction algorithms are different. More detailed report on @tome results is shown in Table S3.

### Supplementary text 2. Author Contributions

All authors participated in conceptualisation, manuscript writing and approved the final version of the manuscript.

**Maria Selifanova —** project supervision, formulation of research goals and aims, data review and validation, drafting original manuscript text, preparation of the illustrations
**Oleg Demianchenko** – genome assembly, annotation plots, identification of ORFanes (stop-codon plots), sequence statistics comparison of protein-coding genes, p-distances and dn/ds comparison of protein-coding genes, data review
**Elizaveta Noskova** – primer design, plots for domain architecture and alignments, reports on general sequence search, clarification of ORFanes boundaries
**Egor Pitikov** – genome assembly and annotation, assembly polishing, phylogenetic trees, CAI calculation, structured segments in sequences
**Denis Skvortsov** – literature search, automatic and manual genome annotation and submission to GeneBank, atp8 identification, genes' alignments and characterisation of protein-coding genes, plots for genome architecture
**Jana Drozd** – domain search, identification of signal peptides, hydropathy profiles of amino acid sequences, some NGS data analysis, functional analysis of ORFanes
**Nika Vatolkina** – distant homology HMM search, structural and functional analysis of ORFanes, literature analysis, search for possible viral origin
**Polina Apel** – primer design, characterisation of protein-coding genes, hydropathy profiles adjustment for alignments
**Ekaterina Kolodyazhnaya** – evolutionary data analysis
**Margarita Ezhova —** library preparations and sequencing, verification PCRs and Sanger sequence reactions.
**Alexander B. Tzetlin —** drafting original manuscript text, specimens identification
**Tatiana V. Neretina —** project supervision, formulation of research goals and aims, sequencing and library preparations, verification PCRs and Sanger sequence reactions, project administration, funding acquisition
**Dmitry A. Knorre —** project supervision, formulation of research goals and aims, drafting original manuscript text, preparation of the illustrations.
